# Single cell sequencing of the small and AT-skewed genome of malaria parasites

**DOI:** 10.1101/2020.02.21.960039

**Authors:** Shiwei Liu, Adam C. Huckaby, Audrey C. Brown, Christopher C. Moore, Ian Burbulis, Michael J. McConnell, Jennifer L. Güler

## Abstract

Single cell genomics is a rapidly advancing field; however, most techniques are designed for mammalian cells. Here, we present a single cell sequencing pipeline for the intracellular parasite, *Plasmodium falciparum*, which harbors a relatively small genome with an extremely skewed base content. Through optimization of a quasi-linear genome amplification method, we achieve better targeting of the parasite genome over contaminants and generate coverage levels that allow detection of relatively small copy number variations on a single cell level. These improvements are important for expanding accessibility of single cell approaches to new organisms and for improving the study of adaptive mechanisms.

## Background

Malaria is a life-threatening disease caused by protozoan *Plasmodium* parasites. *P. falciparum* causes the greatest number of human malaria deaths [1]. The clinical symptoms of malaria occur when parasites invade human erythrocytes and undergo rounds of asexual reproduction by maturing from early forms into late stage parasites and bursting from erythrocytes to begin the cycle again [2]. In this asexual cycle, parasites possess a single haploid genome during the early stages; rapid genome replication in the later stages leads to an average of 16 genome copies [2].

Due to a lack of an effective vaccine, antimalarial drugs are required to treat malaria. However, drug efficacy is threatened by the frequent emergence of resistant populations [3]. Copy number variations (CNVs), or the amplification and deletion of a genomic region, is one of the major sources of genomic variation in *P. falciparum* that contribute to antimalarial resistance [4–15]. Similar to bacteria and viruses [16–18], a high rate of CNVs may initiate genomic changes that contribute to the rapid adaptation of this organism [7, 19]. Despite the importance of CNVs, their dynamics in evolving populations are not well understood.

The majority of CNVs in *P. falciparum* have been identified by analyzing bulk DNA in which the CNVs are present in a substantial fraction of individual parasites in the population due to positive selection [8, 10, 15, 20, 21]. However, many CNVs likely remain undetected because they are presumably either deleterious or offer no advantages for parasite growth or transmission and are therefore present in low frequency [20, 22]. Currently, CNVs can be identified using read-depth analysis of short read sequencing data, which derives an average signal across the population. For this reason, genetic variants must be present in a high frequency (i.e. ∼50%) in the population to be detected [23–25]. Sequencing at very high depth improves the detection of low frequency CNVs, but the sensitivity is limited to large-scale CNVs present in > 5% cells [26–28]. Other analysis methods that rely on the detection of reads that span CNV junctions (i.e. split reads or discordant reads) have improved the sensitivity and specificity of CNV detection [29], but continue to struggle with minor allele detection. This latter method is useful for identifying precise CNV locations, while the read-depth method is required for estimating copy number of CNVs [30]. Because the two methods display distinct sensitivity and specificity for CNV detection, the combination of the two methods improves the accuracy of CNV detection [31].

Recent investigations have analyzed single cells to detect low frequency CNVs within heterogeneous populations [25, 32–36]. This approach provides a significant advantage for detecting rare genetic variants by no longer deriving an average signal from large quantities of cells. However, short read sequencing requires nanogram to microgram quantities of genomic material for library construction, which is many orders of magnitude greater than the genomic content of individual *Plasmodium* cells. Therefore, whole genome amplification (WGA) is required to generate sufficient DNA quantities. Several WGA approaches have been reported and each has advantages and disadvantages for different applications [37–40]; however, most were optimized for mammalian cell analysis [28, 38, 40–51]. Because WGA leads to high levels of variation in read abundance across the genome, CNV analysis in the single cell context is challenging. Previous approaches have been tuned specifically for CNV detection in mammalian genomes, which are generally hundreds of kb to Mb in size [28, 38, 40–51].

To date, the detection of CNVs in single *P. falciparum* parasites using whole genome sequencing has not been achieved. The application of existing WGA methods is complicated by this parasite’s small genome size and extremely imbalanced base composition (23Mb haploid genome with 19.4% GC-content [52]). Each parasite haploid genome contains 25 femtograms of DNA, which is 278-times less than the ∼6400Mb diploid human genome. Therefore, an effective *P. falciparum* WGA method must be both highly sensitive and able to handle the imbalanced base composition. One WGA method, multiple displacement amplification (MDA), has been used to amplify single *P. falciparum* genomes with near complete genome coverage [53, 54]. These studies successfully detected single nucleotide polymorphisms between single parasites but did not report CNV detection, which is possibly disrupted by low genome coverage uniformity [39], the generation of chimeric reads by MDA [55], and the relatively small size of CNVs in *P. falciparum* (broadly <100kb) [20, 22, 56, 57].

Multiple annealing and looping-based amplification cycling (MALBAC) is another WGA method that exhibits improved uniformity over MDA, which is advantageous for detecting CNVs in single cells [27]. MALBAC has the unique feature of quasi-linear pre-amplification, which reduces the bias associated with exponential amplification [27]. However, standard MALBAC is less tolerant to AT-biased genomes, unreliable with low DNA input, and prone to contamination [58–60]. Thus, optimization of this WGA method is necessary for *P. falciparum* genome analysis.

In this study, we developed a single cell sequencing pipeline for *P. falciparum* parasites, which included efficient isolation of single infected erythrocytes, an optimized WGA step inspired by MALBAC, and a sensitive method of assessing sample quality prior to sequencing. We tested our pipeline on erythrocytes infected with laboratory-reared parasites as well as patient-isolated parasites with heavy human genome contamination. Genome amplification using our optimized protocol showed increased genome coverage and better coverage uniformity when compared to standard MALBAC. Furthermore, we have detected CNVs in single cell genomes through the combination of discordant/split reads and read depth analysis methods. Building on these improvements will enable the detection of parasite-to-parasite heterogeneity to clarify the role of genetic variations, such as CNVs, in the adaptation of *P. falciparum*. This study also provides a framework for the optimization of single cell amplification and CNV analysis in other organisms with challenging genomes.

## Methods

### Parasite Culture

We freshly thawed erythrocytic stages of *P. falciparum* (*Dd2*, MRA-150, Malaria Research and Reference Reagent Resource Center, BEI Resources) from frozen stocks and maintained them as previously described [61]. Briefly, parasites were grown in *vitro* at 37°C in solutions of 3% hematocrit (serotype A positive human erythrocytes, Valley Biomedical, Winchester, VA) in RPMI 1640 (Invitrogen, USA) medium containing 24 mM NaHCO_3_ and 25 mM HEPES, and supplemented with 20% human type A positive heat inactivated plasma (Valley Biomedical, Winchester, VA) in sterile, plug-sealed flasks, flushed with 5% O_2_, 5% CO_2_, and 90% N_2_ [7]. We maintained the cultures with media changes every other day and sub-cultured them as necessary to keep parasitemia below 5%. All parasitemia measurements were determined by SYBR green based flow cytometry [62]. Cultures were routinely tested using the LookOut Mycoplasma PCR Detection Kit (Sigma-Aldrich, USA) to confirm negative infection status.

### Clinical Sample Collection

We obtained parasites from an infected patient admitted to the University of Virginia Medical Center with clinical malaria. The patient had a recent history of travel to Sierra Leone, a malaria-endemic country, and *P. falciparum* infection was clinically determined by a positive rapid diagnostic test and peripheral blood smear analysis. We obtained the sample of 1.4% early stage parasites within 24h of phlebotomy, incubated in the conditions described in Parasite Culture for 48 hours and washed the sample 3 times with RPMI 1640 HEPES to decrease levels of white blood cells. In order to fully evaluate our amplification method in the presence of heavy human genome contamination, we did not perform further leukodepletion. We set aside some of the sample for bulk DNA preparation (see *Bulk DNA Extraction*). Using another portion of the sample, we enriched for parasite-infected erythrocytes using SLOPE (Streptolysin-O Percoll) method [63], which increased the parasitemia from 1.4% to 48.5% (**Additional file 1: Figure S1**). We then isolated the single *P. falciparum*-infected erythrocytes using the CellRaft AIR^TM^System (Cell Microsystems, Research Triangle Park, NC) as detailed in *Parasite Staining and Isolation*.

### Bulk DNA Extraction

We lysed asynchronous *P. falciparum*-infected erythrocytes with 0.15% saponin (Sigma-Aldrich, USA) for 5min and washed them with 1x PBS (diluted from 10x PBS Liquid Concentrate, Gibco, USA). We then lysed parasites with 0.1% Sarkosyl Solution (Bioworld, bioPLUS, USA) in the presence of 1mg/ml proteinase K (from *Tritirachium album*, Sigma-Aldrich, USA) overnight at 37°C. We extracted nucleic acids with phenol/chloroform/isoamyl alcohol (25:24:1) pH 8.0 (Sigma-Aldrich, USA) using 2ml light Phase lock Gels (5Prime, USA). Lastly, we precipitated the DNA with ethanol using the standard Maniatis method [64].

### Parasite Staining and Isolation

For late stage parasite samples, we obtained laboratory *Dd2* parasite culture with a starting parasitemia of 1.7% (60% early stage parasites). We separated late stage *P. falciparum*-infected erythrocytes from non-paramagnetic early stages using a LS column containing MACS^®^ microbeads (Miltenyi Biotec, USA, [65]). After elution of bound late stage parasite, the sample exhibited a parasitemia of 80.8% (74.0% late stage parasites, **Additional file 1: Figure S1**). For early stage parasites, we obtained laboratory *Dd2* parasites culture with a starting parasitemia of 3% (46% early stage parasites). We harvested the non-paramagnetic early stages parasites which were present in the flow-through of the LS column containing MACS^®^ microbeads. Next, we enriched the infected erythrocytes using the SLOPE method, which preferentially lysed uninfected erythrocytes [63]. The final parasitemia of enriched early stage parasites was 22.8% (97.0% early stage parasites, **Additional file 1: Figure S1**). To differentiate *P. falciparum*-infected erythrocytes from remaining uninfected erythrocytes or cell debris, we stained the stage specific *P. falciparum*-infected erythrocytes with both SYBR green and MitoTracker Red CMXRos (Invitrogen, USA). We then isolated single *P. falciparum*-infected erythrocytes using the CellRaft AIR^TM^ System (Cell Microsystems, Research Triangle Park, NC). We coated a 100- micron single reservoir array (CytoSort Array and CellRaft AIR user manual, CELL Microsystems) with Cell-Tak Cell and Tissue Adhesive (Corning, USA) following the manufacture’s recommendations. Then, we adhered erythrocytes on to the CytoSort array from a cell suspension of ∼20,000 cells in 3.5mL RPMI 1640 (Invitrogen, USA) with AlbuMAX II Lipid-Rich BSA (Thermo Fisher Scientific, USA) and Hypoxanthine (Sigma-Aldrich, USA). Lastly, we set up the AIR^TM^ System to automatically transfer the manually selected single infected erythrocytes (SYBR+, Mitotracker+) into individual PCR tubes.

### Steps to Limit Contamination

We suspended individual parasite-infected erythrocytes in freshly prepared lysis buffer, overlaid them with one drop (approx. 25μl) of mineral oil (light mineral oil, BioReagent grade for molecular biology, Sigma Aldrich, USA), and stored them at −80°C until WGA. We amplified DNA in a clean positive pressure hood located in a dedicated room, using dedicated reagents and pipettes, and stored them in a dedicated box at −20°C. We wore a new disposable lab coat, gloves and a face mask during reagent preparation, cell lysis, and WGA steps. We decontaminated all surfaces of the clean hood, pipettes, and tube racks with DNAZap (PCR DNA Degradation Solutions, Thermo Fisher Scientific, USA), followed by Cavicide (Metrex Research, Orange, CA), and an 80% ethanol rinse prior to each use. We autoclaved all tubes, tube racks and the waste bin on a dry vacuum cycle for 45min. Finally, we used sealed sterile filter tips, new nuclease-free water (Qiagen, USA) for each experiment, and filtered all salt solutions through a 30mm syringe filter with 0.22μm pore size (Argos Technologies, USA) before use in each experiment.

### Whole Genome Amplification

#### Standard MALBAC

The MALBAC assay was originally designed for human cells [27, 50]. This approach involved making double stranded DNA copies of genomic material using random primers that consist of 5 degenerate bases and 27 bases of common sequence. These linear cycles are followed by exponential amplification of via suppression PCR. Here, we transferred individual cells into sterile thin-wall PCR tubes containing 2.5μl of lysis buffer that yielded a final concentration of 25mM Tris pH 8.8 (Sigma-Aldrich, USA), 10mM NaCl (BAKER ANALYZED A.C.S. Reagent, J.T.Baker, USA), 10mM KCl (ACS reagent, Sigma-Aldrich, USA), 1mM EDTA (molecular biology grade, Promega, USA), 0.1% Triton X-100 (Acros Organics, USA), 1mg/ml Proteinase K (*Tritirachium album,* Sigma-Aldrich, USA). After overlaying one drop of mineral oil, we lysed cells at 50°C for 3h and inactivated the proteinase at 75°C for 20min, then 80°C for 5min before maintaining at 4°C. We added 2.5μl of amplification buffer to each sample to yield a final concentration of 25mM Tris pH 8.8 (Sigma-Aldrich, USA), 10mM (NH_4_)_2_SO_4_ (Molecular biology grade, Sigma-Aldrich, USA), 8mM MgSO_4_ (Fisher BioReagents, Fisher Scientific, Product of India), 10mM KCl (ACS reagent, Sigma-Aldrich, USA), 0.1% Triton X-100 (Acros Organics, USA), 2.5mM dNTP’s (PCR grade, Thermo Fisher Scientific, USA), 1M betaine (PCR Reagent grade, Sigma-Aldrich, USA) and 0.667μM of each random primer (5’GTGAGTGATGGTTGAGGTAGTGTGGAG<u>NNNNN</u>TTT 3’, and 5’GTGAGTGATGGTTGAGGTAGTGTGGAG<u>NNNNN</u>GGG 3’) ordered from Integrated DNA Technologies, USA. To denature DNA, we heated samples to 95°C for 3min and snap-cooled on an ice slush before gently adding 0.5μl of enzyme solution (8,000 U/ml *Bst* DNA Polymerase Large Fragment, New England Biolabs, USA, in 1X amplification buffer) into the aqueous droplet.

We placed the samples into a thermo-cycler (Bio-Rad, USA) holding at 4°C and heated according to the following cycles: 10°C – 45s, 15°C – 45s, 20°C – 45s, 30°C – 45s, 40°C – 45s, 50°C – 45s, 64°C – 10min, 95°C – 20s. The samples were immediately snap-cooled on an ice slush and held for at least 3min to maintain the DNA in a denatured state for the next round of random priming. We added another 0.5μl of enzyme solution and mixed thoroughly with a pipette on ice as above. We placed the samples back into the 4°C thermo-cycler and heated according to the cycles listed above with an additional 58°C step for 1min before once again cooling on an ice slush for 3min. We repeated the addition of enzyme mix (as above) and performed additional rounds of amplification cycles (as above, including the 58°C step). Once completed, we placed the samples on ice and supplemented with cold PCR master mix to yield 50μl with the following concentrations: 0.5μM Common Primer (5’GTGAGTGATGGTTGAGGTAGTGTGGAG3’, Integrated DNA Technologies, USA), 1.0mM dNTPs (PCR grade, Thermo Fisher Scientific, USA), 6.0mM MgCl_2_ (Molecular biology, Sigma-Aldrich, USA), 1X Herculase II Polymerase buffer and 1X Herculase II polymerase (Agilent Technologies, USA). We immediately thermo-cycled samples with the following temperature-time profile: 94°C – 40s, 94°C – 20s, 59°C – 20s, 68°C – 5min, go to step two for several times, and an additional extended at 68°C – 5min, and finally, a hold at 4 °C. For comparison, we used 18/19 linear cycles and 17 exponential cycles for single parasite genomes amplified by the standard MALBAC protocol.

#### Optimized MALBAC

We made the following modifications to standard MALBAC to produce our improved method. **1)** We froze cells at −80°C until usage because freeze-thaw enhanced cell lysis as previously reported [54]; **2)** We removed betaine from the amplification buffer because it improved amplification of GC-rich sequences [66], which are infrequent in *P. falciparum* genomes (**Additional file 2: Table S1**); **3)** We used a single random primer where the GC-content of the degenerate bases were 20% instead of 50% (5’GTGAGTGATGGTTGAGGTAGTGTGGAG<u>NNNNN</u>TTT 3’) at final concentration of 1.2μM; **4)** We reduced the volume of the random priming reaction by added only 0.29μl of 2X amplification buffer to the lysed samples and 0.13μl of enzyme solution to the aqueous droplet each amplification cycle; **5)** We added additional random priming cycles over prior MALBAC studies for a total of 18 (for late stage parasites) or 19 (for early stage parasites) cycles; **6)** We reduced the total volume of exponential amplification from 50μl to 20μl and increased the number of exponential amplification cycles from 15 to 17; **7)** We verified the presence of high molecular weight DNA products in the samples before purifying nucleic acids by Zymo DNA Clean & Concentrator-5 (ZYMO Research).

### Pre-Sequencing Quality Assessment

We assayed 6 distinct genomic loci across different chromosomes to determine variations in copy number following the whole genome amplification step. We included this step, which employs highly sensitive droplet digital PCR (ddPCR, QX200 Droplet Digital PCR system, Bio-Rad, USA), to identify samples that exhibited more even genome coverage prior to short read sequencing. The sequence of primers and probes are described in **Additional file 2: Table S2** [7, 67, 68]. Each ddPCR reaction contained 5μl of DNA (0.3ng/μl for single cell samples), 10μl ddPCR Supermix for Probes (without dUTP), primers and probes with the final concentration in **Additional file 2: Table S2**, and sterile H_2_O to bring the per-reaction volume to 22μl. We prepared droplets with the PCR mixture following the manufacture’s protocol: 95°C – 10 min; 40 cycles of 95°C – 30s, 60°C – 60s, and an infinite hold at 4°C. After thermal cycling, we counted positive droplets using the Bio-Rad QX200 Droplet Reader (Bio-Rad, USA). We analyzed data through QuantaSoft (Bio-Rad, USA). For each gene, a no template control (sterile water, NTC) and a positive control (0.025ng *Dd2* genomic DNA) are included in each ddPCR run. Following ddPCR, we calculated the “uniformity score” using the locus representation of the 6 genes: *seryl tRNA synthetase* (gene-1, PF3D7_0717700)*, heat shock protein 70* (gene-2, PF3D7_0818900)*, dihydrofolate reductase* (gene-3, PF3D7_0417200)*, lactate dehydrogenase* (gene-4, PF3D7_1324900)*, 18S ribosomal RNA* (gene-5, PF3D7_0112300, PF3D7_1148600, PF3D7_1371000), and *multi-drug resistance transporter 1* (*Pfmdr1*, gene-6, PF3D7_0523000) in the amplified DNA sample relative to non-amplified DNA using the following equation:

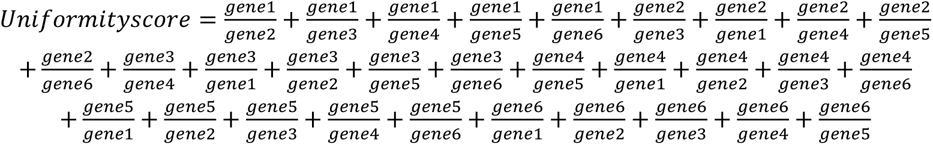

When certain loci were over- or under-represented in the amplified sample, this score increased above the perfect representation of the genome; a uniformity score of 30 indicates that all genes are equally represented. We calculated the locus representation from the absolute copies of a gene measured by ddPCR from 1ng of amplified DNA divided by the absolute copies from 1ng of the bulk DNA control [69]. We only included samples in which all six genes were detected by ddPCR. The relative copy number of the *Pfmdr1*, which was amplified in the *Dd2* parasite line [6], was also used to track the accuracy of amplification. We calculated this value by dividing the ddPCR-derived absolute copies of *Pfmdr1* by the average absolute copies of the 6 assayed loci (including *Pfmdr1*, normalized to a single copy gene). To confirm the efficiency of ddPCR detection as a pre-sequencing quality control step, we determined the strength of association based on the pattern of concordance and discordance between the ddPCR detection and the sequencing depth of the 5 gene targets with Kendall rank correlation (*18S ribosomal RNA* was excluded from correlation analysis due to the mapping of non-unique reads). We then calculated the correlation coefficient (**Additional file 2: Table S3**). When the level of ddPCR detection corresponded to the sequencing depth in at least 3 of the 5 gene targets (a correlation coefficient of >0.6), we regarded the two measurements as correlated.

### Short Read Sequencing

We fragmented MALBAC amplified DNA (>1ng/μL, 50μL) using Covaris M220 Focused Ultrasonicator in microTUBE-50 AFA Fiber Screw-Cap (Covaris, USA) to a target size of 350bp using a treatment time of 150s. We determined the fragment size range of all sheared DNA samples (291bp-476bp) with a Bioanalyzer on HS DNA chips (Agilent Technologies, USA). We used the NEBNext Ultra DNA Library Prep Kit (New England Biolabs, USA) to generate Illumina sequencing libraries from sheared DNA samples. Following adaptor ligation, we applied 3 cycles of PCR enrichment to ensure representation of sequences with both adapters and the size of the final libraries range from 480bp to 655bp. We quantified the proportion of adaptor-ligated DNA using real-time PCR and combined equimolar quantities of each library for sequencing on 4 lanes of an Illumina Nextseq 550 using 150bp paired end cycles. We prepared the sequencing library of clinical bulk DNA as above but sequenced it on an Illumina Miseq using 150bp paired end sequencing.

### Sequencing Analysis

We performed read quality control and sequence alignments essentially as previously described [56] (**Additional file 1: Figure S2A**). Briefly, we removed Illumina adapters and PhiX reads, and trimmed MALBAC common primers from reads with BBDuk tool in BBMap [70]. To identify the source of DNA reads, we randomly subsetted 10,000 reads from each sample by using the reformat tool in BBMap [70] and blasted reads in nucleotide database using BLAST+ remote service. We aligned each fastq file to the hg19 human reference genome and kept the unmapped reads (presumably from *P. falciparum*) for analysis. Then, we aligned each fastq file to the *3D7 P. falciparum* reference genome with Speedseq [71]. We discarded the reads with low-mapping quality score (below 10) and duplicated reads using Samtools [72]. To compare the coverage breadth (the percentage of the genome that has been sequenced at a minimum depth of one mapped read, [73]) between single cell samples, we extracted mappable reads from BAM files using Samtools [72] and randomly downsampled to 300,000 reads using the reformat tool in BBMap [70]. This level is dictated by the sample with the lowest number of mappable reads (ENM, **Additional file 2: Table S4**). We calculated the coverage statistics using Qualimap 2.0 [74] for the genic, intergenic and whole genome regions.

To understand where the primers of MALBAC amplification are annealing to the genome, we overlaid information on the boundaries of genic or intergenic regions with the mapping position of reads containing the MALBAC primer common sequence. To do so, we kept the MALBAC common primers in the sequencing reads, filtered reads and aligned reads as in the above analysis. We subsetted BAM files for genic and intergenic regions using Bedtools, searched for the MALBAC common primer sequence using Samtools, and counted reads with MALBAC common primer using the pileup tool in BBMap (**Additional file 2: Table S5**).

We conducted single cell sequencing analysis following the steps in **Additional file 1: Figure S2B.** We compared the variation of normalized read abundance (log10 ratio) at different bin sizes using boxplot analysis (R version 3.6.1) and determined the bin size of 20 kb using the plateau of decreasing variation of normalized read abundance (log10 ratio) when increasing bin sizes. We then divided the *P. falciparum* genome into non-overlapping 20 kb bins using Bedtools [75]. The normalized read abundance was the mapped reads of each bin divided by the total average reads in each sample. To show the distribution of normalized read abundance along the genome, we constructed circular coverage plots using Circos software and ClicO FS [76, 77]. To assess uniformity of amplification, we calculated the coefficient of variation of normalized read abundance by dividing the standard deviation by the mean and multiplying by 100 [39, 78] and analyzed the equality of coefficients of variation using the R package “cvequality” version 0.2.0 [79]. We employed correlation coefficients to assess amplification reproducibility as previous studies [80]. Due to presence of non-linear correlations between some of the samples, we used Spearman correlation for this analysis. We removed outlier bins if their read abundance was above the highest point of the upper whisker (Q3 + 1.5×interquartile range) or below the lowest point of the lower whisker (Q1-1.5×interquartile range) in each sample. Then, we subsetted remaining bins present in all samples to calculate the correlation coefficient using the R package “Hmisc” version 4.3-0 [81]. We visualized Spearman correlations, histograms and pairwise scatterplots of normalized read abundance using “pairs.panels” in the “psych” R package. We then constructed heatmaps and hierarchical clustering of Spearman correlation coefficient with the “gplots” R package version 3.0.1.1 [82]. Additionally, to estimate the chance of random primer annealing during MALBAC pre-amplification cycles (likely affected by the GC content of genome sequence), we generated all possible 5-base sliding windows with 1 base step-size in the *P. falciparum* genome and calculated the GC-content of the 5-bases windows using Bedtools (**Additional file 2: Table S1**) [75].

We conducted single cell CNV analysis following the steps in **Figure S2C**. To ensure reliable CNV detection, our CNV analysis is limited to the core genome, as defined previously [83]. Specifically, we excluded the telomeric, sub-telomeric regions and hypervariable *var* gene clusters, due to limited mappability of these regions. For discordant/split read analysis, we used LUMPY [84] in Speedseq to detect CNVs with at least two supporting reads in each sample (**Additional file 2: Table S6**). For read-depth analysis, we further filtered the mapped reads using a mapping quality score of 30. We counted the reads in 1kb, 5kb, 8kb, 10kb bins by Bedtools and used Ginkgo to normalize (by dividing the count in each bin by the mean read count across all bins), correct the bin read counts for GC bias, independently segment (using a minimum of 5 bins for each segment), and determine the copy number state in each sample with a predefined ploidy of 1 ([85], **Additional file 2: Table S7**). The quality control steps of Ginkgo were replaced by the coefficient of variation of normalized read count used in this study to assess uniformity in each cell. Lastly, we identified shared CNVs from the two methods when one CNV overlapped with at least 50% of the other CNV and vice versa (50% reciprocal overlap).

## Results

### *Plasmodium falciparum* genomes from single-infected erythrocytes are amplified by MALBAC

Our single cell sequencing pipeline for *P. falciparum* parasites included stage-specific parasite enrichment, isolation of single infected erythrocytes, cell lysis, whole genome amplification, pre-sequencing quality control, whole genome sequencing, and analysis steps (**Figure 1A**). We collected parasites from either an *in vitro*-propagated laboratory line (*Dd2*) or from a blood sample of an infected patient (referred to as ‘laboratory’ and ‘clinical’ parasites, respectively). This allowed us to test the efficiency of our procedures on parasites from different environments with varying amounts of human host DNA contamination. Furthermore, for laboratory samples, we isolated both early (1n) and late (∼16n) stage parasite-infected erythrocytes to evaluate the impact of parasite DNA content on the performance of WGA. For single cell isolation, we used the microscopy-based CellRaft Air system (**Figure 1B**), which has the benefit of low capture volume (minimum: 2μl) and visual confirmation of parasite stages. Following isolation, using the standard MALBAC protocol (termed <u>n</u>on-optimized <u>M</u>ALBAC), we successfully amplified 3 early (ENM) and 4 late stage (LNM) laboratory samples. We also applied a version of MALBAC that we optimized for the small AT-rich *P. falciparum* genome (termed <u>o</u>ptimized <u>M</u>ALBAC) to 42 <u>e</u>arly (EOM) and 20 late stage (LOM) laboratory samples as well as 4 <u>c</u>linical samples (COM) (**Additional file 2: Table S8**). Compared to standard MALBAC, our optimized protocol had a lower reaction volume, more amplification cycles, and a modified pre-amplification random primer (see *Methods* for more details). Using this method, we successfully amplified 43% of the early and 90% of the late stage laboratory samples and 100% of the clinical samples (see post-amplification yields in **Additional file 2: Tables S8 and S9**).

**Figure 1.**
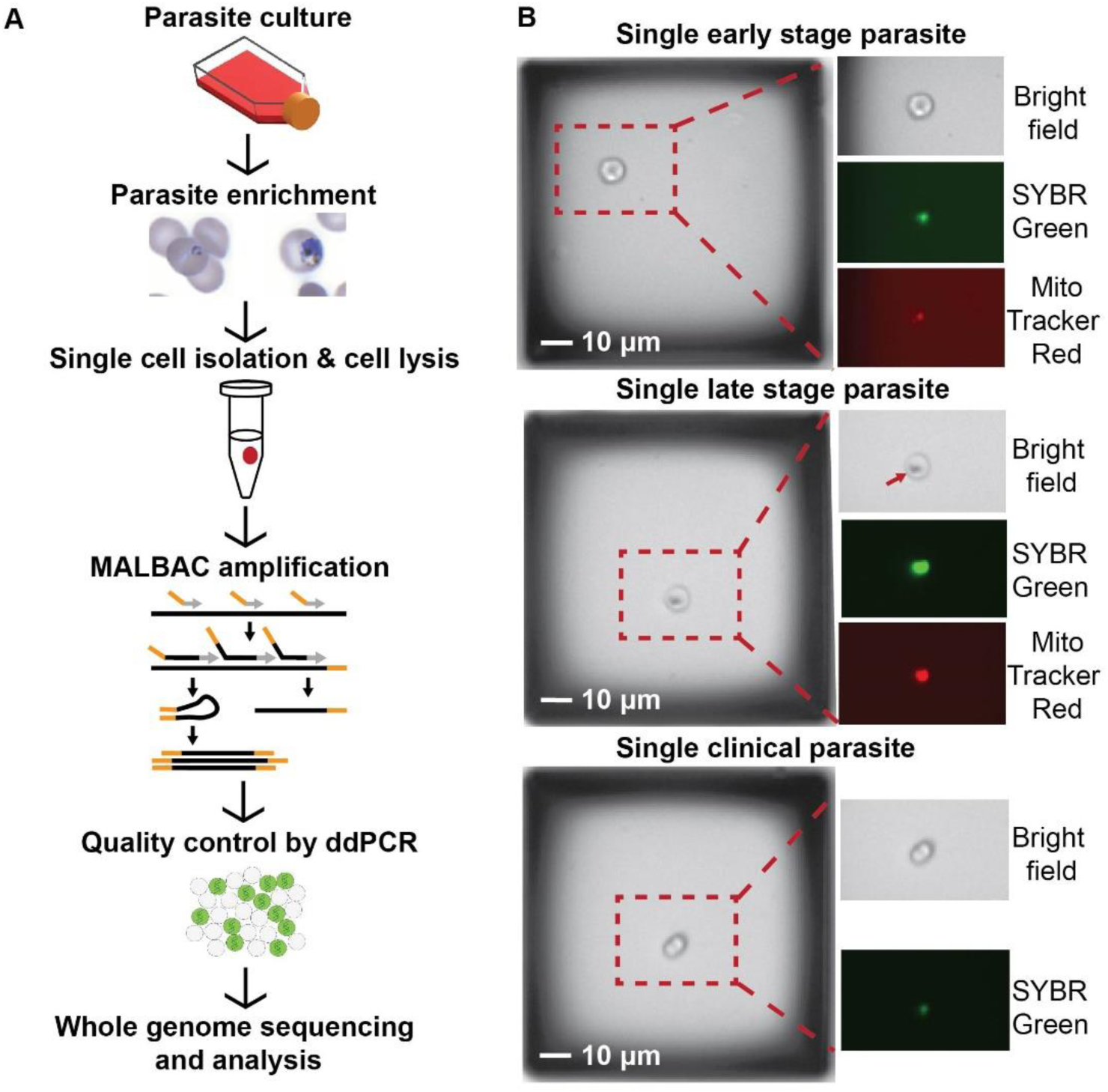
**Single *P. falciparum*-infected erythrocytes are isolated, amplified, and sequenced. A**. **Experimental workflow.** Parasites are grown *in vitro* in human erythrocytes or isolated from infected patients. In order to limit the number of uninfected erythrocytes in the sample, infected cells are enriched using column and gradient-based methods (see *Methods*). Individual early-stage (left image) and late-stage (right image) parasite-infected erythrocytes were automatically isolated into PCR tubes using the CellRaft AIR System (Cell Microsystems, see panel B). All cells were lysed by combining a freeze–thaw step with detergent treatment prior to MALBAC amplification. MALBAC uses a combination of common (orange) and degenerate (grey) primers to amplify the genome. The quality of amplified genomes was assessed prior to library preparation and sequencing using droplet digital (dd) PCR; DNA is partitioned into individual droplets to accurately measure gene copies. Suitable samples were Illumina sequenced and analyzed as detailed in **Additional file 1: Figure S2**. **B. Parasite stage visualization on the CellRaft AIR System using microscopy** (10X magnification). Enriched early and late stage parasite*-*infected erythrocytes at low density were seeded into microwells to yield only a single cell per well (left image of each group), and identified with SYBR green and Mitotracker Red staining (indicates parasite DNA and mitochondrion, respectively). Early stage parasites exhibited lower fluorescence due to their smaller size and late stage parasites had noticeable dark spots (arrow) due to the accumulation of hemozoin pigment. Scale bar represents 10μm.

### A novel pre-sequencing quality control step identifies samples with more even genome amplification

We assessed the quality of WGA products from single cells using droplet digital PCR (ddPCR) to measure the copy number of single and multi-copy genes dispersed across the *P. falciparum* genome (6 genes in total including *Pfmdr1*, which is present at ∼3 copies in the *Dd2* laboratory parasite line). Using this sensitive quantitative method, along with calculation of a “uniformity score” which reflects both locus dropout and over-amplification, we were able to select genomes that had been more evenly amplified; a low uniformity score and accurate copy number values indicated a genome that has been amplified with less bias (see *Methods* for details on score calculation and **Additional file 2: Table S10** for primary data). This quality control step was important to reduce unnecessary sequencing costs during single cell studies.

When we analyzed differences between successfully amplified samples by optimized MALBAC (17 EOM samples and 14 LOM samples processed for ddPCR evaluation) and non-optimized MALBAC (3 ENM and 4 LNM samples), we found that samples amplified with the optimized protocol were more evenly covered than those from the standard method (**Table 1**). Based on the results of ddPCR detection, we selected a subset of 13 EOM and 10 LOM samples for sequencing (**Additional file 2: Table S8**). Overall, selected samples had lower average uniformity scores (i.e. 248 and 1012 for selected and unselected EOMs, respectively, see **Table 1**). For clinical parasite samples, 3 out of 4 COM samples showed a lack of ddPCR detection on at least one parasite gene; thus, we were not able to calculate a uniformity score for these samples and the amplification of clinical genomes was likely more skewed than laboratory samples (**Table 1**).

**Table 1.**
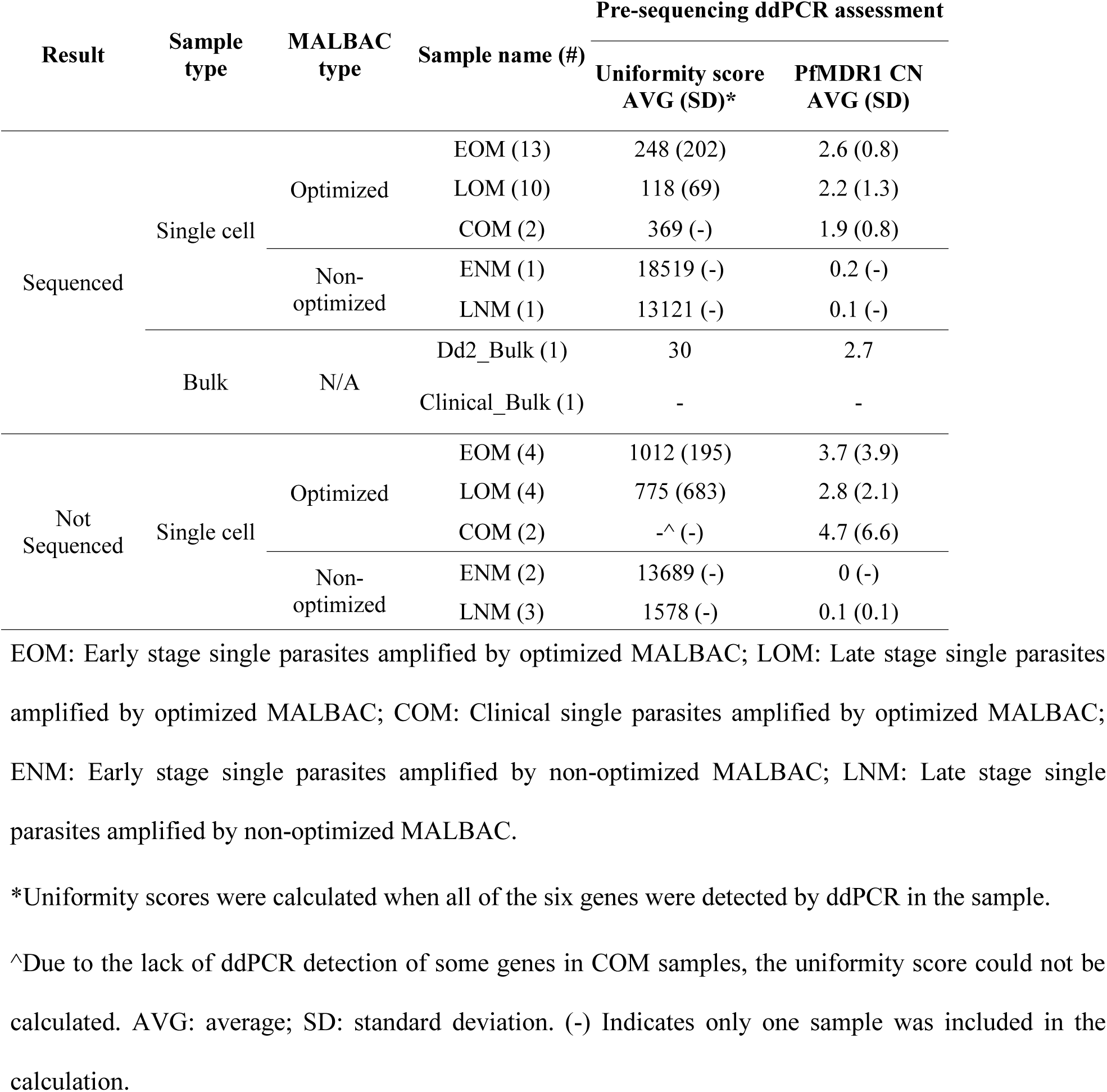
Pre-sequencing quality control by droplet digital PCR

Both standard and optimized MALBAC-amplified parasite genomes were short read sequenced alongside a matched bulk DNA control (**Table 1**). To confirm the efficiency of ddPCR detection as a pre-sequencing quality control step, we calculated the correlation between ddPCR quantification and the sequencing depth at these specific loci. We found that the ddPCR-derived gene copy concentration was correlated with sequencing coverage of the corresponding genes in many samples (**Additional file 2: Table S3**, 17 out of 28 samples are correlated, Kendal rank correlation coefficient >= 0.6), confirming the validity of using ddPCR detection as a quality control step prior to sequencing.

### Optimized MALBAC limits contamination of single cell samples

After read quality control steps, we mapped the reads to the *P. falciparum 3D7* reference genome (see *Methods* and **Additional file 1: Figure S2** for details). We first assessed the proportion of contaminating reads in our samples; NCBI Blast results showed that the majority of non-*P. falciparum* reads were of human origin (**Figure 2A**). The proportions of human reads in 6 out of 13 EOM samples (1.1%-6.9%) and 8 out of 10 LOM samples (1.4%-6.1%) were lower than that in the control bulk sample (7.4%, **Figure 2A**). Conversely, the proportion of human reads in ENM and LNM samples were much higher (81.8% and 18.9%, respectively). As shown in other studies [86, 87], our clinical bulk DNA (81.9%) contained a much higher level of human contamination than the laboratory *Dd2* bulk DNA (7.4%). However, we found that the proportion of the human contaminating DNA in the two single cell COM samples was considerably lower (58.8% and 65.5%). The second most common source of contaminating reads was from bacteria such as *Staphylococcus* and *Cutibacterium*. The ENM sample exhibited a ∼10- fold increase in the proportion of bacterial reads over averaged EOM samples (8.2% versus 0.8%, respectively) whereas the LNM samples showed the same proportion of bacterial reads as the averaged LOM samples (0.2%). These results indicated that the optimized MALBAC protocol reduced the amplification bias towards contaminating human and bacterial genomes.

**Figure 2.**
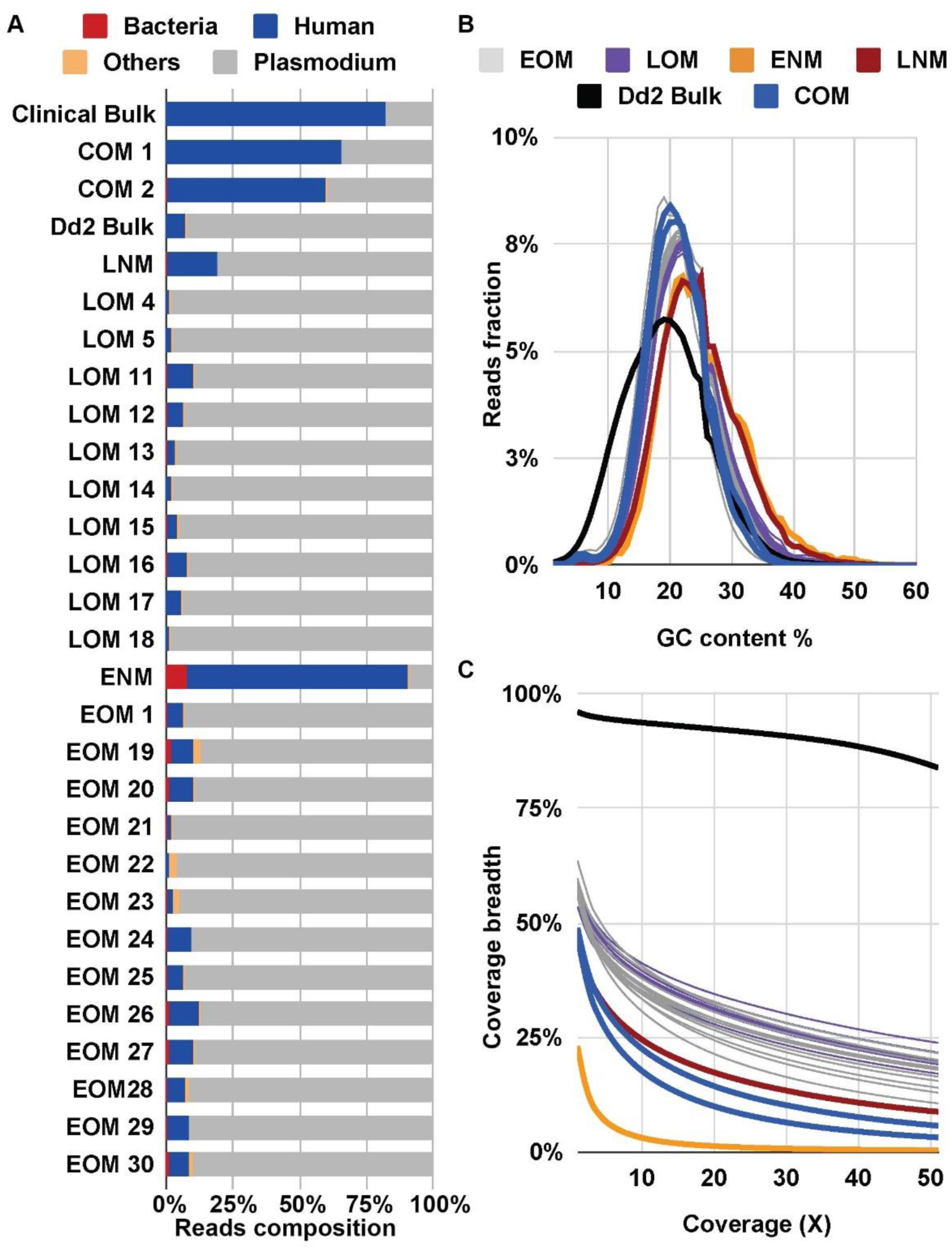
**Sequencing statistics show benefits of optimized MALBAC. A. Contribution of reads based on organism type**. A subset of 10,000 reads from each sample were randomly selected for BLAST to identify sources of DNA. Color representation: bacteria (red); human (blue); other organisms (orange); *Plasmodium* (grey). **B**. **GC-content of *P. falciparum* mapped reads.** GC-content of reads was calculated by Qualimap with default parameters. Color representation: EOM (grey): Early stage single parasites amplified by optimized MALBAC; LOM (purple): Late stage single parasites amplified by optimized MALBAC; ENM (orange): Early stage single parasites amplified by non-optimized MALBAC; LNM (dark red): Late stage single parasites amplified by non-optimized MALBAC; *Dd2* bulk genomic DNA (black); COM samples (blue): Clinical single parasites amplified by optimized MALBAC. Clinical Bulk genomic DNA is not shown here due to <1% of the genome being covered by at least one read. **C. Fraction of *P. falciparum* genome covered by >1 read.** The fraction of the genome was calculated by Qualimap with default parameters. Color representations are the same as described in panel B.

### Optimized MALBAC reduces amplification bias of single cell samples

To further assess the optimized MALBAC protocol, we evaluated GC-bias at several steps of our pipeline (i.e. WGA, library preparation, and the sequencing platform itself). Analysis of the bulk genome control (without WGA) indicated that there was little GC-bias introduced by the library preparation, sequencing, or genome alignment steps; the GC-content of mapped reads from bulk sequencing data is 18.9% (**Table 2**), which was in line with the GC-content (19.4%) of the reference genome [52]. We then compared values from single cell samples to those from the appropriate bulk control to evaluate the GC-bias caused by MALBAC amplification (**Figure 2B**). The average GC-content of all EOM (21.4%), LOM (22.4%), and COM (20.7%) samples was within 1-3.5% of the bulk controls from laboratory and clinical samples (18.9% and 19.7%, respectively, **Table 2**). However, the average GC-content of ENM and LNM samples was 6.1% and 5.4% greater than that of the bulk control; this results is consistent with the high GC preference of the standard protocol [38, 60]. ENM and LNM samples also showed a greater proportion of mapped reads with high GC-content (>30%) than EOM, LOM, and bulk DNA samples (**Figure 2B**).

**Table 2.**
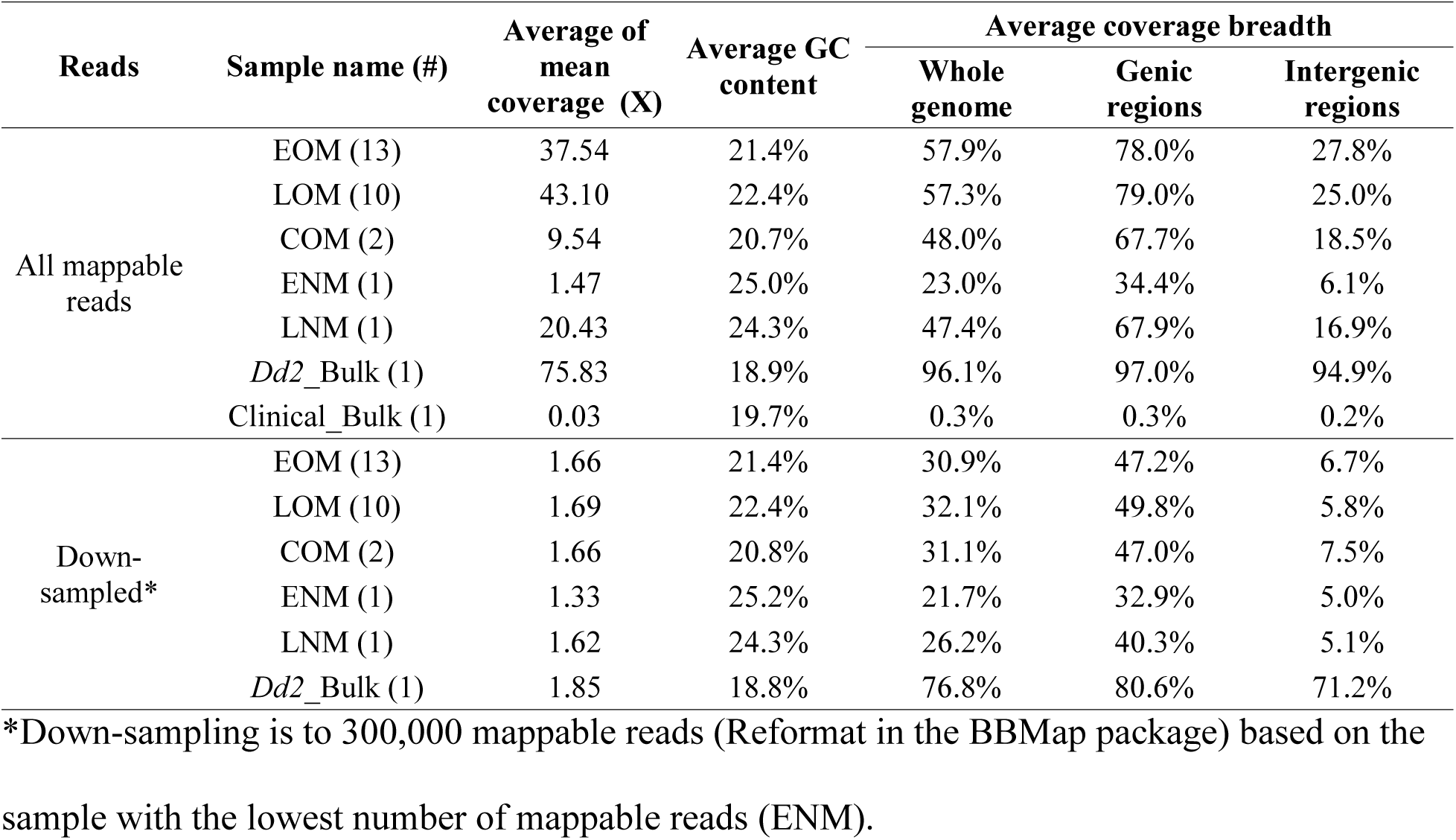
Average GC-content and coverage breadth of sequenced samples

Since GC-bias during the amplification step can limit which areas of the genome are sequenced, we assessed whether the optimization of MALBAC improved genome coverage. The coverage breadth of single cell samples increased by 34.9% in early stage samples (**Figure 2C**, orange-ENM to grey-EOM lines) and by 9.9% for late stage samples following optimization (**Figure 2C**, red-LNM to purple-LOM lines, see values in **Table 2**). Even when we randomly down-sampled reads to the same number per sample (300,000), EOM and LOM samples continued to show improved coverage breadth over ENM and LOM samples (**Table 2**). Even though optimized MALBAC showed less bias towards GC-rich sequences, it was still problematic for highly AT-rich and repetitive intergenic regions (mean of 13.6% GC-content, [52]). The fraction of intergenic regions covered by reads was only 27.8% for EOM samples and 25.0% for LOM samples on average. When we excluded intergenic regions, the fraction of genic regions of the genome covered by at least one read reached an average of 78.0% and 79.0% for EOM and LOM samples (**Table 2**). Conversely, the coverage of intergenic and genic regions was significantly lower for the non-optimized samples. Coverage of the *P. falciparum* genome in the clinical bulk sample was very low due to heavy contamination with human reads (0.3% of the genome was covered by at least one read). This was much lower than that from the 2 COM samples (an average of 48%, **Figure 2C** and **Table 2**).

### Optimized MALBAC improves uniformity of single cell genomes

To investigate the uniformity of read abundance distributed over the *P. falciparum* genome, we divided the reference genome into 20kb bins and plotted the read abundance in these bins over the 14 chromosomes (**Figure 3A, Additional file 1: Figure S3 and S4A**). We selected a 20kb bin size based on its relatively low coverage variation compared to smaller bin sizes and similar coverage variation as the larger bin sizes (**Additional file 1: Figure S5**). To quantitatively measure this variation, we normalized the read abundance per bin in each sample by dividing the raw read counts with the mean read counts per 20kb bin (**Figure 3B, Additional file 1: Figure S3C**). Here, the bulk control displayed the smallest range of read abundance for outlier bins (blue circles) and lowest interquartile range (IQR) value of non-outlier bins (black box, **Figure 3B, Additional file 1: Figure S3C**), indicating less bin-to-bin variation in read abundance. Both EOM and LOM samples exhibited a smaller range of normalized read abundance in outlier bins than ENM and LNM samples (**Figure 3B, Additional file 1: Figure S3C**). In addition, the read abundance variation of COM samples was similar to EOM or LOM samples (**Figure 3B, Additional file 1: Figure S4B**). Finally, due to the extremely low coverage of the clinical bulk sample, the read abundance variation was much higher than all other samples (**Figure 3B, Additional file 1: Figure S4B**).

**Figure 3.**
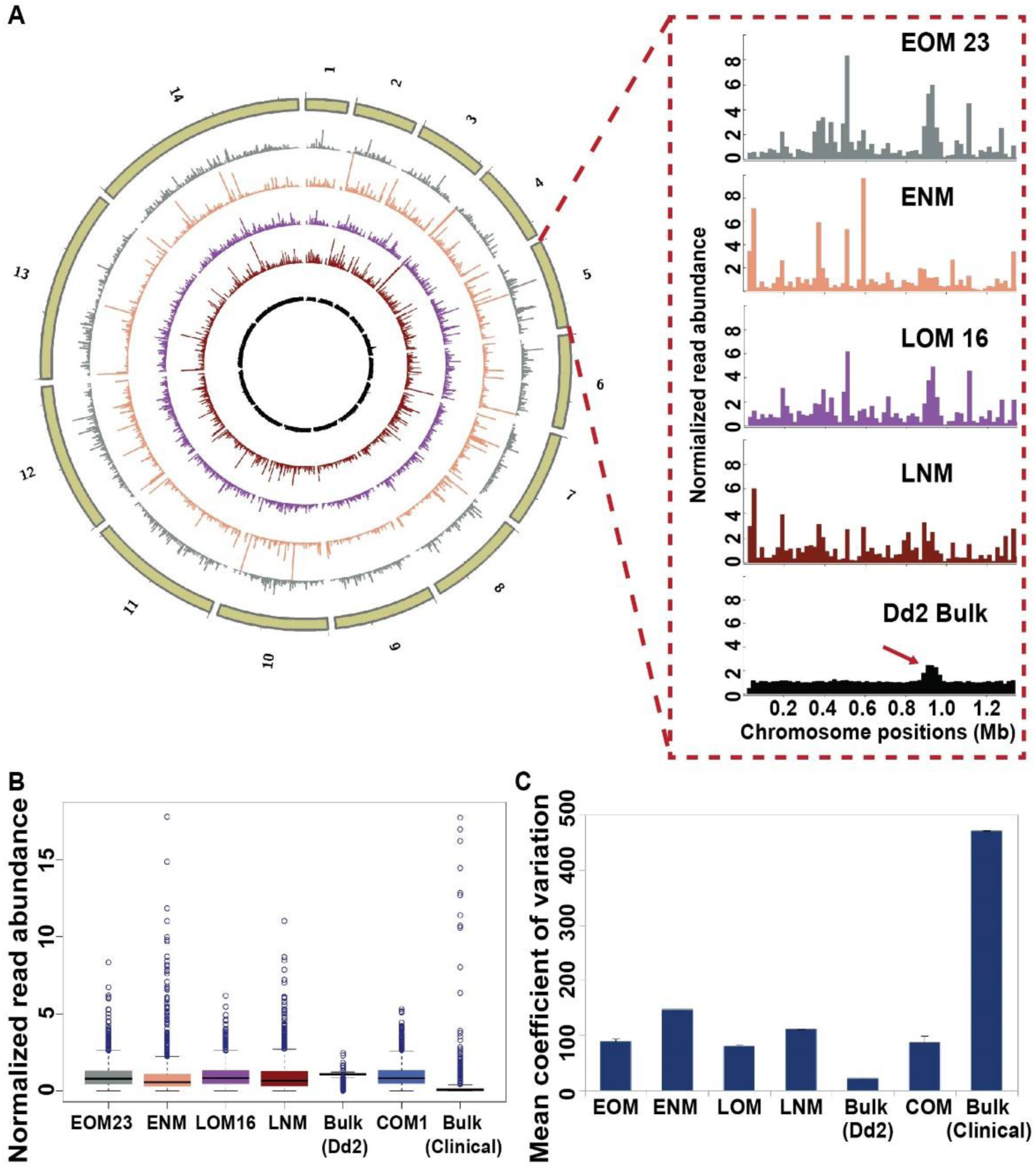
**Samples amplified by optimized MALBAC display improved uniformity of read abundance. A. Normalized read abundance across the genome**. The reference genome was divided into 20kb bins and read counts in each bin were normalized by the mean read count in each sample. The circles of the plot represent (from outside to inside): chromosomes 1 to 14 (tan); one EOM sample (#23, grey); one ENM sample (#3, orange); one LOM sample (#16, purple); one LNM sample (#2, dark red); *Dd2* bulk genomic DNA (black). The zoomed panel shows the read distribution across chromosome 5, which contains a known CNV (*Pfmdr1* amplification, arrow on *Dd2* bulk sample). **B. Distribution of normalized read abundance values for all bins.** The boxes were drawn from Q1 (25^th^ percentiles) to Q3 (75^th^ percentiles) with a horizontal line drawn in the middle to denote the median of normalized read abundance for each sample. Outliers, above the highest point of the upper whisker (Q3 + 1.5×IQR) or below the lowest point of the lower whisker (Q1-1.5×IQR), are depicted with circles. One sample from each type is represented (see all samples in **Additional file 1: Figure S3C**). **C. Coefficient of variation of normalized read abundance.** The average and SD (error bars) coefficient of variation for all samples from each type is represented (EOM: 13 samples; ENM: 1 sample; LOM: 10 samples; LNM: 1 sample; Dd2 Bulk: 1 sample; COM: 2 samples; Clinical Bulk: 1 sample). See *Methods* for calculation.

We then calculated the mean coefficient of variation (CV) for read abundance in the different sample types (**Table 3, Figure 3C, Additional file 2: Table S11**). Following normalization for coverage, the CV from the ENM sample was significantly higher compared to the CV of each EOM sample (147% versus a mean of 89%, respectively, pairwise p value < 0.01, **Additional file 2: Table S12**). Similarly, the LNM-CV was significantly higher compared to the CV of each LOM sample (111% versus a mean of 79%, respectively, pairwise p value <0.01, **Additional file 2: Table S12**). These data showed improvement in levels of read uniformity across the genome when using optimized MALBAC over the standard protocol. In support of this finding, the CV value of COM samples was similar to EOM and LOM samples (**Table 3, Figure 3C**).

**Table 3.** Coefficient variation of normalized read abundance in each sample type

| Sample name | Mean Coefficient of Variation (CV) | SD |
| --- | --- | --- |
| *Dd2* Bulk (1) | 22 | - |
| ENM (1) | 147 | - |
| EOM (13) | 89 | 4 |
| LNM (1) | 111 | - |
| LOM (10) | 79 | 2 |
| COM (2) | 87 | 12 |
| Clinical Bulk (1) | 472 | - |

SD, standard deviation.

### Optimized MALBAC exhibits reproducible coverage of single cell genomes

To better assess the amplification patterns across the genomes, we compared the distribution of binned normalized reads from single cell samples to the bulk control using a correlation test (as performed in other single cell studies [38, 88]). This analysis revealed that amplification patterns of optimized EOM and LOM samples were slightly correlated with the bulk control (Spearman correlation coefficient of 0.27 and 0.25, respectively, **Additional file 2: Table S13**), while the non-optimized samples were not correlated (ENM: 0.05 and LNM: 0.07) (**Figure 4A**). This result indicated that the parasite genome was better represented by single cell samples amplified by optimized MALBAC. To quantify the reproducibility of read distribution between single cell samples amplified by MALBAC, we compared their Spearman correlation coefficients. The read abundance across all single cell samples was highly correlated; two individual EOM or LOM samples had a correlation coefficient of 0.83 and 0.88 respectively (**Figure 4B**). When we expanded our analysis to calculate the correlation of binned normalized reads between all 26 sequenced samples (**Additional file 2: Table S13**) and hierarchically clustered the Spearman correlation coefficient matrix between these samples, all 23 optimized single cell samples (EOM and LOM) clustered with a mean Spearman correlation coefficient of 0.84 (**Figure 4C**). In addition, the two COM samples were correlated with each other (Spearman correlation coefficient of 0.84) (**Additional file 1: Figure S4**C). This correlation indicated high reproducibility of normalized read distribution across the genomes that were amplified by optimized MALBAC. Within the large cluster, two LOM samples (LOM12 and LOM13) displayed the highest correlation (0.94, **Figure 4C**).

**Figure 4.**
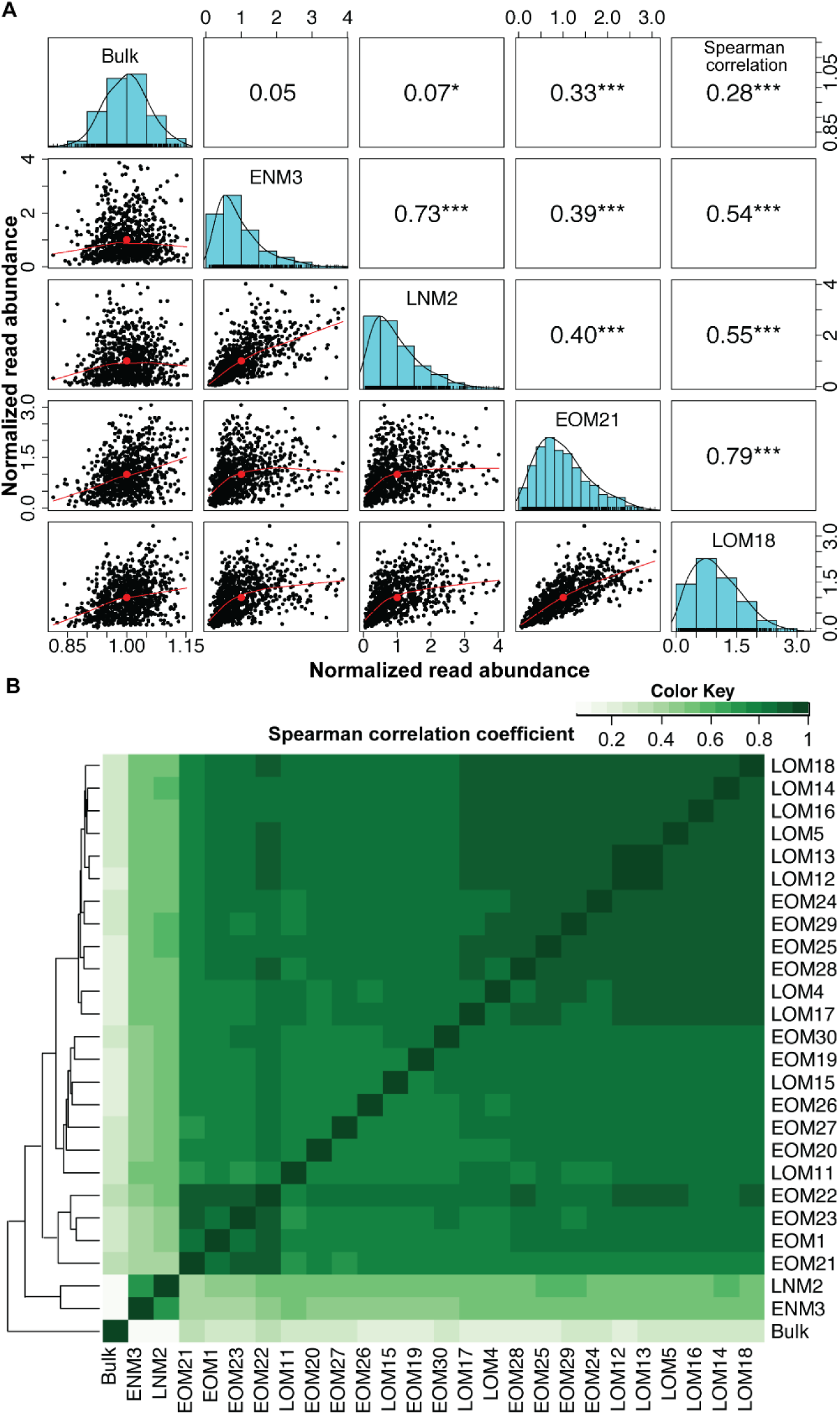
**Correlations show reproducibility of amplification pattern by optimized MALBAC. A.** Paired panels for 5X5 matrices represent Spearman correlation, histogram and pairwise scatterplot among the normalized read abundance of the *Dd2* Bulk, ENM, LNM, and one of each EOM and LOM samples. Outlier bins were removed prior to this analysis (see *Methods* for outlier identification). The Spearman correlation coefficients of each pair are listed above the diagonal, and stars indicate the p-value at levels of 0.1 (no star), 0.05 (*), 0.01 (**), and 0.001 (***). The histograms on the diagonal shows the distribution of normalized read abundance in each sample. The bivariate scatter plots, below the diagonal, depict the fitted line through locally smoothed regression and correlation ellipses (an ellipse around the mean with the axis length reflecting one standard deviation of the x and y variables). **B. Spearman correlation coefficients between sequenced samples.** The hierarchical clustering heatmap was generated using Spearman correlation coefficients of normalized read abundance. The color scale indicates the degree of correlation (white, correlation= 0; green, correlation > 0).

### Reproducible coverage with lower variation is the main benefit of MALBAC over MDA-based amplification of single cell genomes

We performed a brief comparison between our data and that from a MDA-based study because this is the only other method that has been used to amplify single *Plasmodium* genomes ([54], **Additional file 1: Figure S6**). This study sorted individual infected erythrocytes with high (H), medium (M) and low (L) DNA content corresponding to late, mid, and early stage parasites, applied MDA-based WGA to single erythrocytes, and sequenced the DNA products. The authors measured a similar amplification success rate in early (L) stage samples as our study (MDA: 50% by DNA yield, MALBAC: 43% by DNA yield) yet slightly improved success rates for late (H) stage samples (MDA: 100%, MALBAC: 90%, **Additional file 2: Table S8 and S9**). In light of experimental differences between the two studies (**Additional file 2: Table S14**), we analyzed data from the twelve MDA samples using our exact analysis pipeline and parameters (six MDA-H and three of each MDA-M and -L samples) and confined our comparison of the data to a few metrics: 1) coefficient of variation of read abundance, 2) coverage breadth, and 3) correlation between samples (see below).

While MALBAC-amplified genomes exhibited a consistent amplification pattern (**Additional file 1: Figure S3A and S3B**), the MDA-amplified genomes showed substantially more variation across cells (**Additional file 1: Figure S6A**). We also detected higher variation in normalized read abundance in the MDA-H samples (compared to MDA-L and -M samples, **Additional file 1: Figure S6B**), which was not consistent with the report that the MDA method amplifies high DNA content better than parasites with lower DNA content [54]. Even though the bulk DNA controls used in both studies showed similar CVs (24% versus 22%), the MDA-amplified samples displayed a higher CV than MALBAC-amplified single cell samples regardless of the parasite stage (a mean of 186% versus 85%, respectively, **Table 3, Additional file 2: Table S11 and S15**). Additionally, the correlation between MDA-amplified cells (mean correlation coefficient: 0.20; **Additional file 2: Table S17, Additional file 1: Figure S6D**) was much lower than that between our optimized MALBAC-amplified cells (mean correlation coefficient: 0.84; **Additional file 2: Table S13, Figure 4C**). As expected based on MALBAC’s limited coverage of intergenic regions (**Table 2**), MDA amplified samples displayed a higher coverage breadth cross the genome, especially in the intergenic regions (**Additional file 2: Table S16**).

### Copy number variation analysis is achievable in MALBAC-amplified single cell genomes

To detect CNVs with confidence, we employed both discordant/split read detection and read-depth based methods with strict parameters. We used LUMPY to detect paired reads that span CNV breakpoints or have unexpected distances/orientations (requiring a minimum of 2 supporting reads). We also used a single cell CNV analysis tool, Ginkgo, to segment the genome based on read depth across bins of multiple sizes and determine copy number of segments (requiring a minimum of 5 consecutive bins). We regarded the CNVs detected by the two methods the same if one CNV overlapped with at least half of the other CNV and vice versa (50% reciprocal overlap). Using this approach, we first identified a “true set” of CNVs from the bulk *Dd2* DNA sample (**Table 4**, 3 CNVs on 3 different chromosomes). One of the true CNVs was identified previously in this parasite line (the large *Pfmdr1* amplification on chromosome 5, [6]); another true CNV occurs in an area of the genome that is reported to commonly rearrange in laboratory parasites ([89], the *Pf11-1* amplification of chromosome 10).

**Table 4.** True CNVs detected in the Dd2 bulk genome

| Name | Chr. | Start Pos. | Size (bp) | Type | Support read* |  | Start Pos. | Size (bp) | Copy number detected by Ginkgo** in different bin sizes |  |  |  | Mappability^ |
| --- | --- | --- | --- | --- | --- | --- | --- | --- | --- | --- | --- | --- | --- |
|  |  |  |  |  | Discordant read | Split read |  |  | 1kb | 5kb | 8kb | 10kb |  |
| *Pfmdr1* | 5 | 888316 | 81935 | DUP | 53 | 0 | 888000 | 82000 | 2 | 2 | Nd | Nd | 1 |
| *Pf11-1* | 10 | 1524527 | 18472 | DUP | 29 | 1 | 1520000 | 28000 | 4 | 5 | NA | NA | 0.86 |
| *Pf332* | 11 | 1956623 | 8719 | DUP | 0 | 8 | 1953000 | 13000 | 4 | NA | NA | NA | 0.92 |

\*Detected by LUMPY based on discordant/split read detection, minimum number of supporting reads is 2.
\*\*For Ginkgo analysis, the minimum bin number of segmentation is 5.
^For comparison, the mean mappability of the core genome is 0.99 and the mean mappability telomere/subtelomere regions including *var* gene clusters is 0.65.
DUP, duplication; NA, not applicable because the target CNVs will not be detected as the bin size ( $\geq 5 \times$ bin size) is larger than the size of the target CNVs. Nd, not detected.

With a set of true CNVs in hand, we assessed our ability to detect these CNVs in the single cell samples amplified by MALBAC and explored parameters that impacted their detection. As expected, each CNV detection method exhibited differences in ability to identify the true CNVs, which is likely due to a number of factors including CNV size, genomic neighborhood, and sequencing depth [31]. For example, using discordant/split read analysis, we were able to readily identify the *Pf11-1* amplification in the majority of samples (21 of 25 samples, **Additional file 2: Table S18**). This method was less successful in identifying the *Pfmdr1* amplification (only 3 optimized MALBAC samples in total, **Additional file 2: Table S18).** For read-depth analysis, the success of true CNV detection was heavily dependent on the bin size (**Additional file 2: Table S18**). If we selected the lowest bin size (1kb) in which it was possible to detect the smallest of the true CNVs (13kb), we were able to readily identify the *Pfmdr1* amplification in all samples (**Additional file 2: Table S18**). As we increased the bin size, the number samples with *Pfmdr1* detection decreased, only optimized MALBAC samples were represented, and the copy number estimate in single cells approached the bulk control (**Additional file 2: Table S7 and S18**). The other two true CNVs were only detected at the 1kb bin size in a minority of samples (6 total, **Additional file 2: Table S18).**

When we assessed true CNVs that overlapped between the two methods, we were able to detect at least one CNV in a total of 5 single cells (3 EOM and 2 LOM samples out of 25 total cells, **Table 5**). In one sample, EOM 23, the *Pfmdr1* amplification was detected in bin sizes of up to 10kb at a copy number similar to the bulk control (∼5 copies, **Table 5**). Besides the CNVs conserved with the *Dd2* bulk sample, we also detected unique CNVs that were only identified in the single cell samples. In general, most of the CNVs detected by both discordant/split read and read depth analyses were spread across all but one chromosome (including 1-8, 10-14), predominantly confined to optimized MALBAC samples, and were only detected at 1kb read depth bin sizes (**Additional file 2: Table S19**).

**Table 5.**
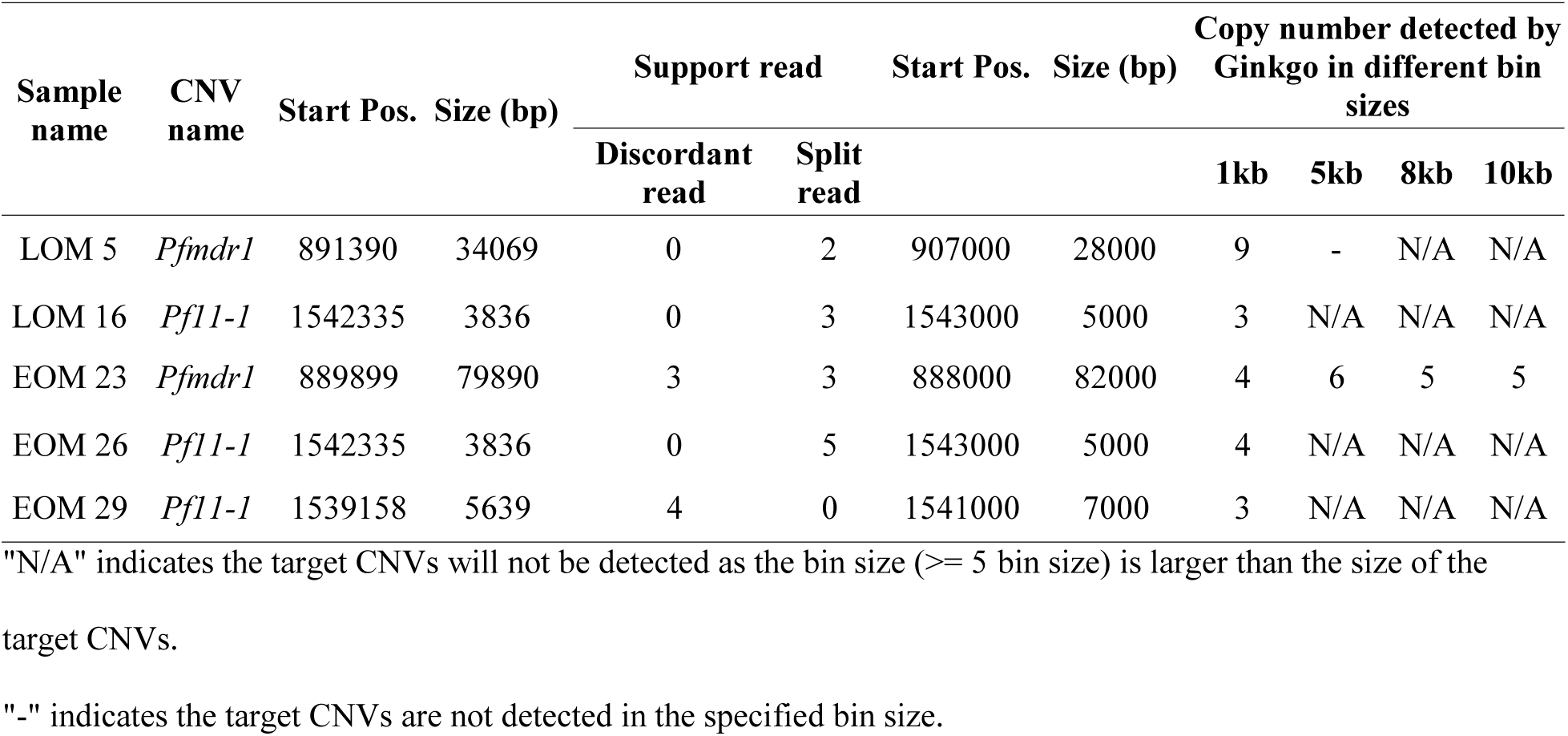
True CNVs detected in single cells

## Discussion

This study is the first to optimize the standard MALBAC protocol for single cell sequencing of a genome with extreme GC-content (*P. falciparum*: 19.4%). We showed that this optimized method can reliably amplify early stage parasite genomes, which contain <30 femtograms of DNA per sample. Single cell experiments are innately very sensitive to contaminating DNA from other organisms and we detected a lower proportion of human and bacteria DNA in MALBAC-amplified samples, which improved overall coverage of the *P. falciparum* genome. Furthermore, we showed that this method reduced GC-bias to increase the breadth and uniformity of genome amplification; these improvements contributed to the detection of true CNVs in single parasite genomes.

### MALBAC Volume and Cycles

MALBAC amplification has been used in studies of human cells, where each single genome harbors a picogram level of DNA [27, 50]. In this study, we successfully improved the sensitivity of the MALBAC method to amplify a femtogram level of DNA from single *P. falciparum* parasites. Reducing the total reaction volume (from 50μl to 20μl) and increasing the number of amplification cycles (pre-amplification: from 5 to 19-20; exponential: from 15 to 17) was likely responsible for this improvement in sensitivity. It was essential to combine these two changes; the lower sample volume and decreased starting material reduced the overall DNA yield and therefore, we increased the number of amplification cycles to generate enough material for sequencing. Additional benefits of these modifications included less contaminating DNA introduced by reagents and reduced costs due to the lower reagent requirement. Importantly, these simple steps can be applied to the MALBAC amplification of small genomes or genomes with skewed GC-content from other organisms such as bacteria [90]. For example, studies of *Mycoplasma capricolum* (GC-poor) [91]*, Rickettsia prowasekii* (GC-poor) [92], and *Borrelia burgdorferi* (GC-poor) [93], *Entamoeba histolytica* (GC-poor) [94]*, Micrococcus luteus* (GC-rich) [95] could be improved using this method.

### Primers and Coverage Bias

The modification of the primer was essential for the successful amplification of the AT-rich *P. falciparum* genome. This change was meant to prevent the preferential amplification of GC-rich sequences as observed for human and rat single cell genomes [38, 60]. We increased coverage breadth of *P. falciparum* genic regions (a mean of 21.7% GC-content) from as low as <40% to ∼80% (ENM versus EOM and LOM samples, **Table 2**) by specifically altering the base content of the degenerate 5-mer of MALBAC pre-amplification primer from 50% to 20% GC-content. The initial priming step is crucial for whole genome amplification and controlling this step can limit amplification bias [96]. Indeed, 5-mers with ∼20% GC-content across the *P. falciparum* genome are 2- and 6-fold more common than those with 40% and 60% GC-content, respectively (**Additional file 2: Table S1**). This difference indicated that annealing of the optimized MALBAC primer based on the degenerate bases was more specific for the parasite’s genome than the standard MALBAC primer. Interestingly, during this study we observed a preferential amplification of genic over intergenic regions (**Table 2**), which may be explained by lower percentage of 5-mers with 20% GC-content in intergenic regions than in genic regions (**Additional file 2: Table S1**). Furthermore, when we searched for reads that contained the MALBAC common sequence (see *Methods* and **Additional file 2: Table S5**) to identify WGA binding sites across the *P. falciparum* genome, we found that binding sites were predominantly located in the genic regions (**Additional file 2: Table S5**); this result indicated that there was an issue with primer annealing in intergenic regions, which may be caused by a high predicted rate of DNA secondary structure formation across these regions of the *P. falciparum* genome [56]. The polymerase used in the MALBAC linear amplification steps (*Bst* large fragment) exhibits strand displacement activity, which presumably allows resolution of secondary structure [97, 98]. However, a longer extension time may be required for amplification of repetitive DNA sequence, either during linear or exponential steps.

### Parasite and Contaminating Genomes

The standard MALBAC method is reported to display the most favorable ratio of parasite DNA amplification over human DNA when compared to other common WGA methods [99]. Our optimization of MALBAC further improved this ratio. The improved sensitivity of optimized MALBAC through reducing reaction volume and increasing cycle numbers not only enhanced the amplification of the small parasite genome, but also improved the sensitivity to amplify contaminating non-parasite DNA. Nevertheless, when comparing the two MALBAC protocols, the optimized method yielded a greatly reduced proportion of contaminating DNA (ENM and LNM: 13.6% vs EOM and LOM: 6.9% of total reads, **Figure 2A**). We speculate that this decrease was once again due to our modification of the GC-content of the degenerate bases of the primer; this limited the preferential amplification by standard MALBAC of contaminating DNA with higher GC content, improving the representation of parasite DNA in the overall WGA product.

The major contaminating DNA source that we detected in our samples was from humans (**Figure 2A**). This was not surprising given that, in our experimental system, the culture and host environments are rich in human DNA [86, 87, 100]. It is also possible that human DNA was introduced during the single cell isolation or WGA steps [59]. The former situation is a larger issue for clinical parasite isolates due to the abundance of white blood cells that contribute to extracellular DNA when they decay outside of the host [101]. Indeed, we observed more human DNA in clinical bulk and single cell samples (an increase of ∼11-fold over laboratory-derived *Dd2* bulk and single cell samples, respectively). The massive level of contamination in the clinical bulk sample and limited coverage of the parasite genome (0.3%) was exacerbated by 1) the omission of a leukodepletion step that is routinely employed to limit host cell contamination [102–104] and 2) the lower overall sequencing output of that particular run (**Additional file 2: Table S4**).

The second most common source of contaminating DNA was bacteria (**Figure 2A**). Since this contaminant was absent in the bulk DNA control and increased in early stage parasite samples (representing an average of 0.8% of EOM reads compared to 0.2% for LOM samples), we predict that bacterial material was introduced during single cell isolation and WGA steps. Although we took precautions to limit this occurrence (see *Methods*), environmental cells and DNA could have been introduced during parasite isolation using the open microscopy chamber of the CellRaft AIR System. In addition, other potential sources include the molecular biology grade water [105–107] or WGA reagents [108–111]. Reducing the reaction volume could further reduce this source of contamination.

### Early and Late Stage Parasites

Depending on when a novel CNV arises (i.e. early or late in replication), each parasite stage holds advantages for its detection. If the CNV arises in the first round of replication and is copied into most of the genomes of a late stage parasite, having multiple genomes will be advantageous for detection. If the CNV arises later in replication, it will be present in only few of the genomes; therefore, averaging across the genomes, as with bulk analysis, will limit its detection. Since only one haploid genome is present in an early stage parasite, the sensitivity for detecting rare/unique CNVs within parasite populations will be enhanced in this situation.

For this reason, staging of parasites in this study was important. We performed stage-specific enrichment before single cell isolation and confirmed that the majority of parasites were the desired stage using flow cytometry (see *Methods*, **Additional file 1: Figure S1**, 97% for early stage enrichment and 74% for late stage enrichment). Furthermore, during selection of cells by microscopy (before automated collection by the IsoRaft instrument), we confirmed the expected fluorescence intensities for each stage; early stage parasites had a significantly smaller genome and mitochondrion size compared to late state (as in **Figure 1B**). However, differences in preparation of samples may have impacted our parasite stage comparisons. While all late stage samples were isolated, lysed and amplified in the same batch under the same conditions, early stage samples processed in three separate batches (**Additional file 2: Table S11**). Despite conserved methods and good concordance in CV between all samples (**Additional file 2: Table S11**), minor differences could have contributed to variations in the amplification steps.

Differences in our genome analysis results from optimized MALBAC samples provided further confidence that the parasites were of the expected stage. Firstly, late stage parasites showed a higher WGA success rate than early stage parasites (90% versus 43%, **Additional file 2: Table S9**). This result was explained simply by the presence of extra genomes in the late stage samples (∼16n versus 1n) and was consistent with a previous study that used MDA-based amplification methods [54]. Late stage parasites also displayed better uniformity of read abundance (**Table 3**), indicating less amplification bias because fewer regions were missed when more genomes were present. Additionally, there were fewer contaminating reads found in late stage parasites than early stage parasites overall (5.1% versus 8.6%). Once again, this was likely due to a higher ratio of parasite DNA to contaminating DNA in the late stage samples.

Despite these differences, we observed similar coverage breadth and Spearman correlation coefficients of read abundance for both early and late stage MALBAC-amplified parasites (**Table 2 and Additional file 2: Table S13**). This was contrary to the MDA study in single *P. falciparum* parasites that found a higher breadth of genome coverage from the late stage parasites [54]. Our findings confirmed that the pattern of amplification across the genome was determined by the binding of the optimized MALBAC primers and not the parasite developmental form.

### Amplification Reproducibility and CNV Analysis

The high level of amplification reproducibility (i.e. the same regions are over- and under-amplified across multiple genomes), that we and others have observed with MALBAC, is especially advantageous for CNV detection because amplification bias can be normalized across cells (as has been successfully performed for human cells [27, 112]). However, cross-sample normalization is not possible in our study due to the use of a single laboratory parasite line that includes known CNVs (*Dd2*). Instead, we lowered our false positive rate by combining a read-depth based tool (Ginkgo) with a split/discordant read-based method (LUMPY) to detect CNVs in our single cell samples. Using this approach, we identified at least one true CNV in a minority of single cell genomes (*Pfmdr1* or *Pf11.1* amplifications were detected in 5 of 25 samples, **Table 5**). However, for read-depth analysis, these calls were confined to the 1kb bin size; this observation may be explained by a number of possibilities, including those that are both biological and artifacts of our methods. For example, the parameters of Ginkgo may be limiting CNV detection at larger bin sizes (requires a minimum 5 bins to call a CNV) or because random noise is higher at this bin size, the false positive rate is higher and therefore the random chance for overlap with LUMPY calls is increased. From a biological perspective, however, there may be an abundance of small CNVs as has been observed by genomic studies on this parasite [22]. Ultimately, additional validation with larger sample sizes will be required to determine the answer.

Importantly, as we increased the bin size, the uniformity of read count improves (**Figure S5**) and impacts our ability to identify CNVs (i.e. the *Pfmdr1* amplification is found in fewer single cell genomes and the copy number estimate approaches that of the bulk control, **Table S18 and S7**). Thus, we assert that we can accurately detect relatively large CNVs (>50kb) in single parasite samples using larger bin sizes (>=10kb). This is an advancement in single cell genomics for two reasons: 1) we have identified a ∼82kb CNV in single cell genomes that is below the current resolution of CNV detection from single cell genomes amplified with common WGA methods (hundreds of kb to Mb) [27, 28, 46, 51, 60, 113–115] and 2) our sensitivity for CNV detection will improve greatly when we add cross-sample normalization to our analysis pipeline. This step will be possible when we expand our studies in number and parasite diversity; the inclusion of parasite lines with different CNV profiles along the genome will greatly facilitate the removal of reproducible amplification bias and increase the reproducible detection of conserved and unique CNVs of all sizes.

### Limitations, Scope, and Future Efforts

One limitation in our comparison between standard and optimized MALBAC-amplified samples was the sequencing of only a single standard MALBAC sample from each parasite stage. However, we evaluated a total of 7 independent non-optimized samples (3 ENM and 4 LNM) and detected multiple instances of allelic dropout, could not calculate the uniformity score for 4 of 7 samples, and detected heavy skewing of the copy number of a known CNV (**Table 1 and Additional file 2: Table S10**). These results indicated biased coverage and high levels of contaminating DNA in these samples, which made sequencing of these samples futile.

Additionally, since our goal in this study was to evaluate amplification bias, we did not perform SNP analysis on samples to address accuracy of the MALBAC method. Other studies showed that the WGA-induced single nucleotide error rate with MALBAC was higher than that for MDA [27, 59, 116]. This was likely due to the use of error-prone large fragment *Bst* polymerase in MALBAC pre-amplification cycles compared to the use of phi29 DNA polymerase with proofreading activity in MDA.

While it is notable that we can successfully amplify a small, base-skewed genome and generate coverage levels that allow the detection of relatively small CNVs on a single cell level, we recognize that improvements can be made to our CNV analysis pipeline. As mentioned above, future studies will include the use of cross-sample normalization to increase our accuracy of CNV detection. Additionally, it will be important to further explore the genomic features associated with amplification bias; for example, the annealing location of common sequences of MALBAC primers and the location of secondary structure in the *P. falciparum* genome could impact amplification steps [117]. In this case, if associations are identified, we can further normalize for these features in a similar manner as we currently do so for GC content difference across bins. Any improvements in the coverage of intergenic regions and uniformity will also impact CNV identification through increased detection of discordant/split reads and more accurate read-depth calling in these regions.

## Conclusions

Our modifications of reaction volume, cycle number, and GC-content of degenerate primers will expand the use of MALBAC-based approaches to organisms not previously accessible because of small genome size or skewed base content. Furthermore, these changes can reduce amplification of undesired contaminating genomes in a sample. The reproducible nature of this WGA method, combined with new genome analysis tools, will reduce the effect of amplification bias when conducting large scale single cell analysis and enhance our ability to explore genetic heterogeneity. Thus, we expect this approach to broadly improve study of mechanisms of genetic adaptation in a variety of organisms.

## List of abbreviations

MALBAC: Multiple annealing and looping-based amplification cycling
CNVs: Copy number variations
WGA: Whole genome amplification
MDA: Multiple displacement amplification
PfMDR1: Plasmodium falciparum multidrug resistance 1
ddPCR: Droplet digital PCR
NA: Not applicable
SD: Standard deviation
EOM: Early stage single parasites amplified by optimized MALBAC
LOM: Late stage single parasites amplified by optimized MALBAC
COM: Clinical single parasites amplified by optimized MALBAC
ENM: Early stage single parasites amplified by non-optimized MALBAC
LNM: Late stage single parasites amplified by non-optimized MALBAC
IQR: Interquartile range
CV: Coefficient of variation
SLOPE: Streptolysin-O Percoll

## Declarations

### Ethical Approval and Wavier for Informed Consent

The University of Virginia Institutional Review Board for Health Sciences Research provided ethical approval for clinical samples used in this study (IRB-HSR protocol #21081). We handled all samples in accordance with approved protocols and in agreement with ethical standards of the Declaration of Helsinki. The University of Virginia Institutional Review Board for Health Sciences Research provided a wavier for informed consent because our study design met the following criteria: the research involved minimal risk to subjects, the waiver does not adversely affect the rights and welfare of subjects, and the research could not practicably be carried out without the waiver.

### Availability of data and materials

The raw sequence files generated and analyzed during the current study are available in the Sequence Read Archive (SRA) under the BioProject ID PRJNA607987, BioSamples SAMN14159290-SAMN14159318. The datasets for the uniformity and reproducibility analysis of MDA-based amplification on parasite DNA from single infected erythrocytes are available in the NCBI short read archive under the accession PRJNA385321[54].

### Authors’ contributions

MJM and JLG conceived of the project. SL, ACH, and JLG designed the experiments. IB and MJM provided access to essential protocols and equipment (CellRaft AIR System) at the start of the project. ACB and CCM procured and processed clinical samples from the University of Virginia Medical Center. SL conducted all of the experiments. SL analyzed the data, with support from ACH and JLG. SL and JLG wrote the manuscript. ACH, ACB, CCM, IB, and MJM edited the manuscript. All authors critically reviewed and approved the manuscript.

## Acknowledgements

Our thanks to Michelle Warthan for parasite culturing support, AnhThu Nguyen (Biology Genomics Core, University of Virginia) for sequencing support, and all members of Dr. Jennifer Guler’s laboratory, as well as the laboratories of Dr. William Petri Jr and Barbara Mann at University of Virginia, for their helpful discussion and insight.

