## Additional file 1 for "Single cell sequencing of the small and AT-skewed genome of malaria parasites"

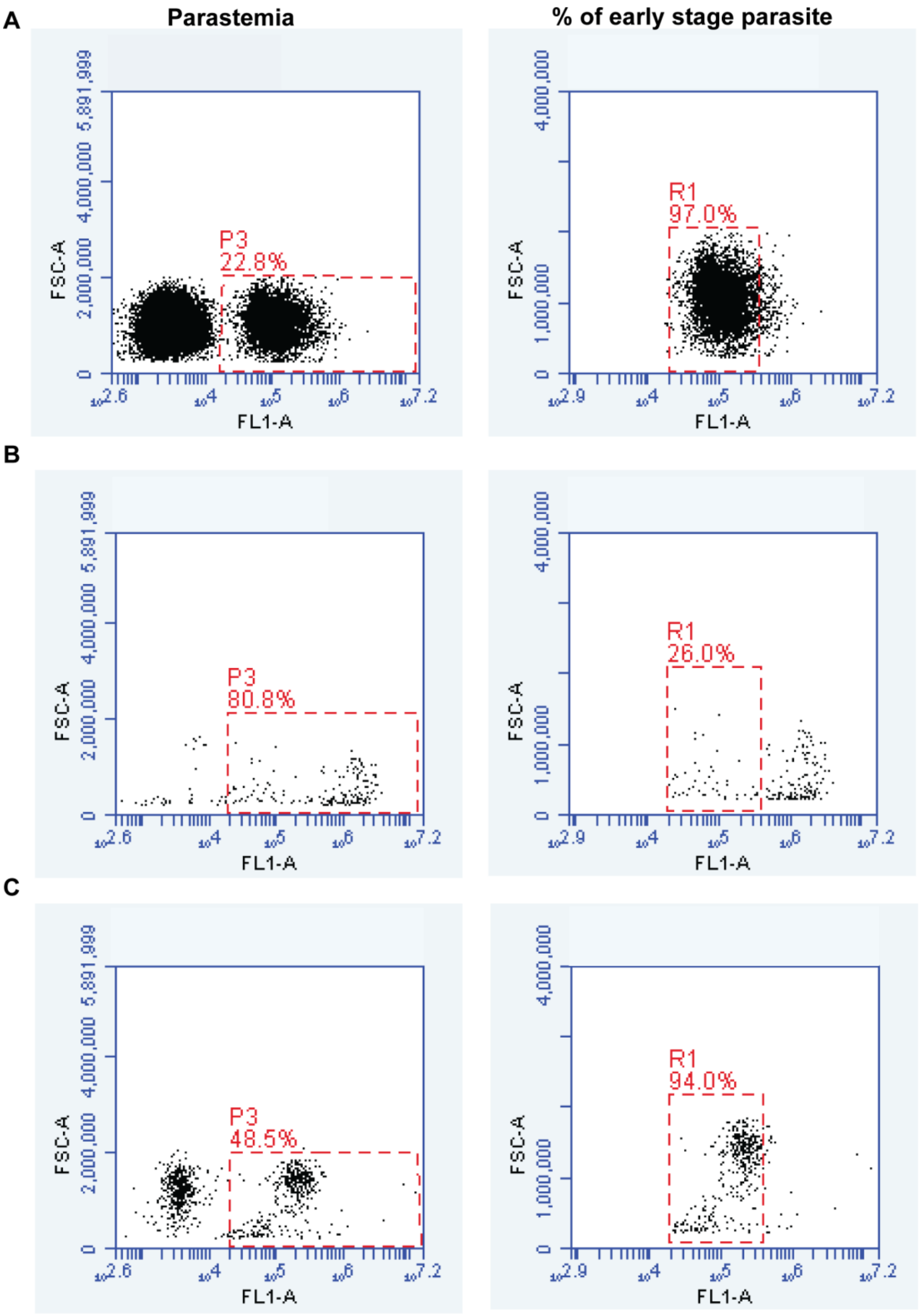


**Figure S1. Confirmation of staging for enriched parasite samples.** Parasitemia (proportion of infected erythrocytes; P3 gate, left plots) and proportion of early stage parasites (R1 gate, right plots) is shown for each sample. **A. Early stage laboratory parasite samples.** Early stage parasites were enriched by harvesting the flow-through of the MACS column, which contained parasite stages that have not accumulated hemozoin, a paramagnetic crystal, along with uninfected cells. The SLOPE method was used to deplete the sample of uninfected cells to yield a high parasitemia (22.8%), predominantly early-staged population (97.0%) for single cell isolation. The flow cytometry run collected 30,485 total events. **B. Late stage laboratory parasite samples.** Late stage parasites were enriched by collecting the bound fraction from the MACS column, which contained parasite stages with high levels of paramagnetic hemozoin. This high parasitemia (80.8%), predominantly late stage population (74%), was used for single cell isolation. Late-stage parasites were further selected by higher fluorescence (due to increased DNA content and mitochondrial size) on the CellRaft microscope during isolation (see **Figure 1A**). The flow cytometry run collected 1357 total events. **C. Early stage clinical parasite samples**. Whole blood collected from a *P. falciparum*-infected patient was stored in sodium citrate for 23 hours in the hospital and incubated in RPMI for 48 hours in the laboratory (see *Methods* for details). The SLOPE method was used to deplete the sample of uninfected cells to yield a high parasitemia (48.5%), predominantly early-staged population (94.0%, confirmed by microscopy) for single cell isolation. The flow cytometry run collected 1314 total events.

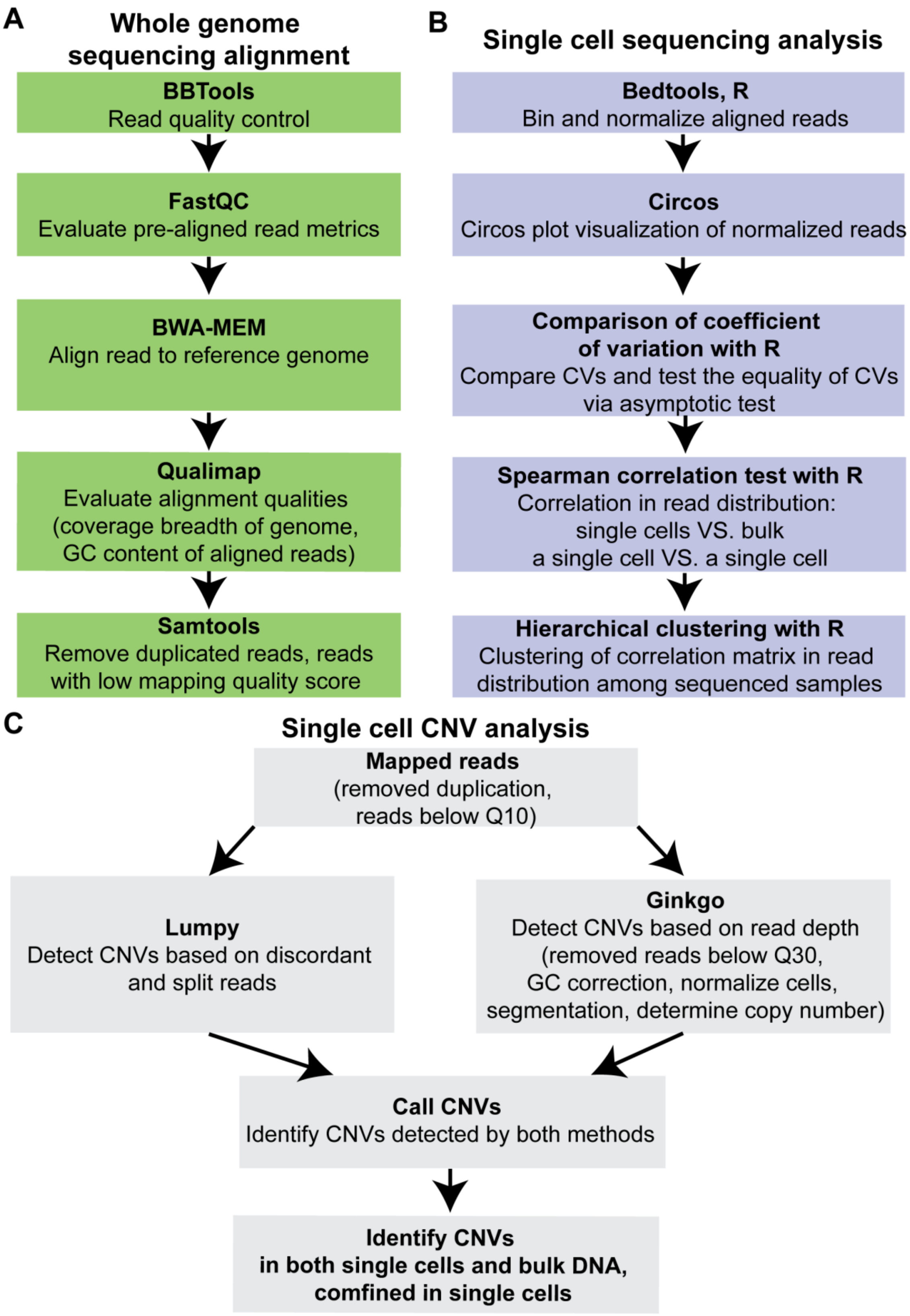


**Figure S2. Bioinformatic analysis of sequencing reads. A. Whole genome sequencing analysis and alignment.** Alignment of whole genome sequencing reads started with BBTools to remove low quality bases, adapter sequences, trim the common sequence of MALBAC primer and verify correct pairing of reads. The resulting “clean” paired reads were evaluated by FastQC for overrepresented sequences, per base read qualities, and read length distributions. After passing read quality control, BWA-MEM was used to align “clean” paired reads to the *3d7 Plasmodium falciparum* reference genome. Qualimap was then used to evaluate the alignments for the fraction of the genome covered by at least one read, GC-content of aligned reads, and mapping quality. Duplicated reads and reads with mapping quality score below 10 were removed for downstream analysis by Samtools. **B. Single cell sequencing analysis steps.** The reference genome was divided into 20kb bins by Bedtools. Read counts was calculated in every 20kb bins and normalized by the mean read count in each sample with Bedtools and R. Circos was utilized to visualize the distribution of normalized read counts over 14 chromosomes. To compare variance of read counts in each sample, coefficient of variations in read counts were calculated with R and the equality of coefficient of variations was tested via Feltz and Miller’s (1996) asymptotic test. Correlation coefficients between all combinations of pairs of sequenced samples were calculated via Spearman correlation test on normalized read counts in 20kb bins with R. Clustering of Spearman correlation matrix of normalized read counts in 20kb bins was conducted among sequenced samples with R. **C. CNV analysis steps.** Mapped reads were filtered by removing duplicated reads and applying a cutoff for mapping quality score of 10. LUMPY required at least two supporting reads for CNV detection. After mapped reads were further filtered by a mapping quality score of 30, Ginkgo was used to detect CNVs based on read depth across 1kb, 5kb, 8kb, 10kb bins; steps included normalization, GC content correction of read counts, independent segmentation, and copy number determination for each sample. CNVs were called by identifying those that are shared between the LUMPY Ginkgo (see *Methods* for details).

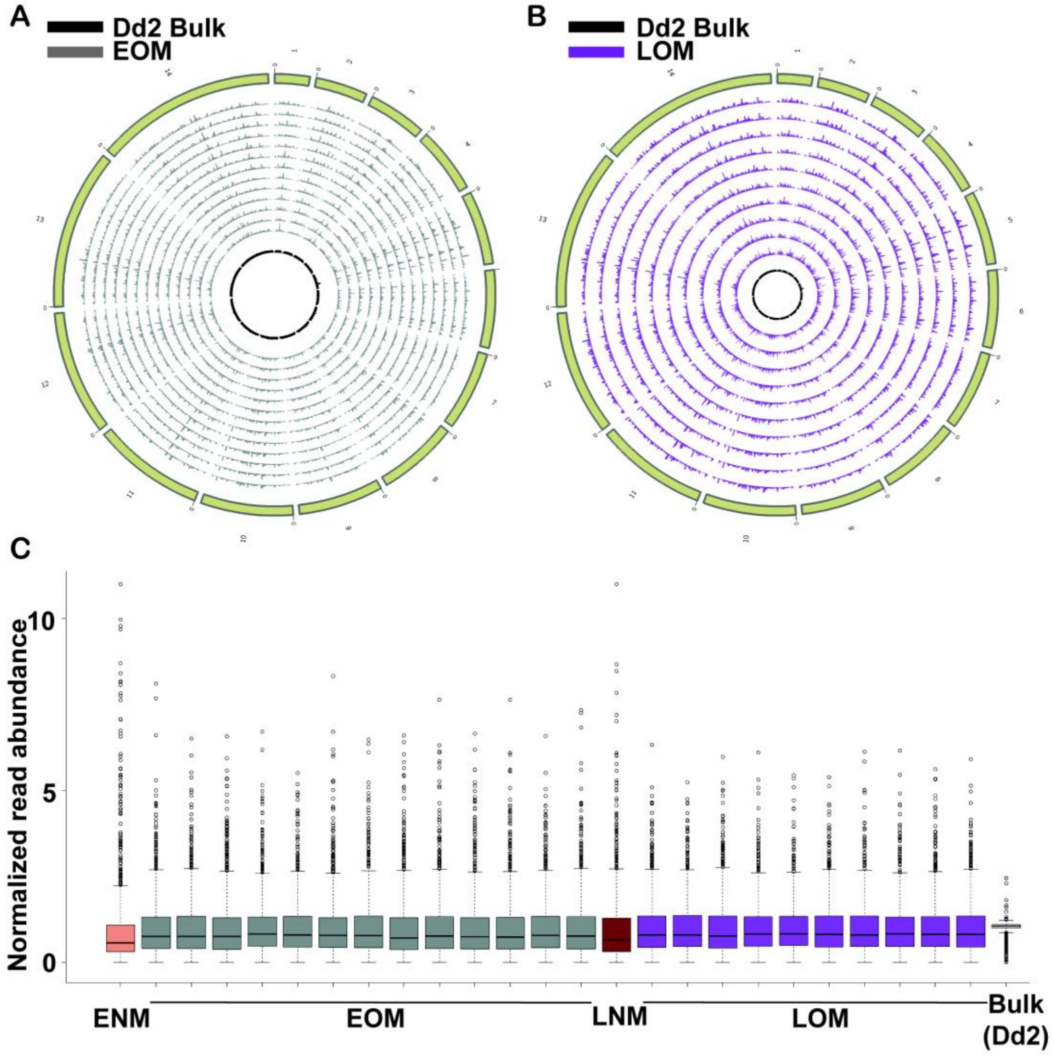


**Figure S3. Uniformity of read abundance across the whole genome of all EOM and LOM samples. A & B. Normalized read abundance across the genome**. Distribution of read abundance in 20kb bins in all EOM (A) and LOM (B) samples is shown. Read counts in each bin were normalized by the mean read count over the whole genome for each sample. The circles from outside to the inside represent: chromosomes 1:14 (tan); 13 EOM samples (grey) or 10 LOM samples (purple); Bulk genomic DNA (black). **C. Distribution of normalized read abundance values for all bins**. Normalized read abundance per 20kb bins with outliers (circles) is represented.

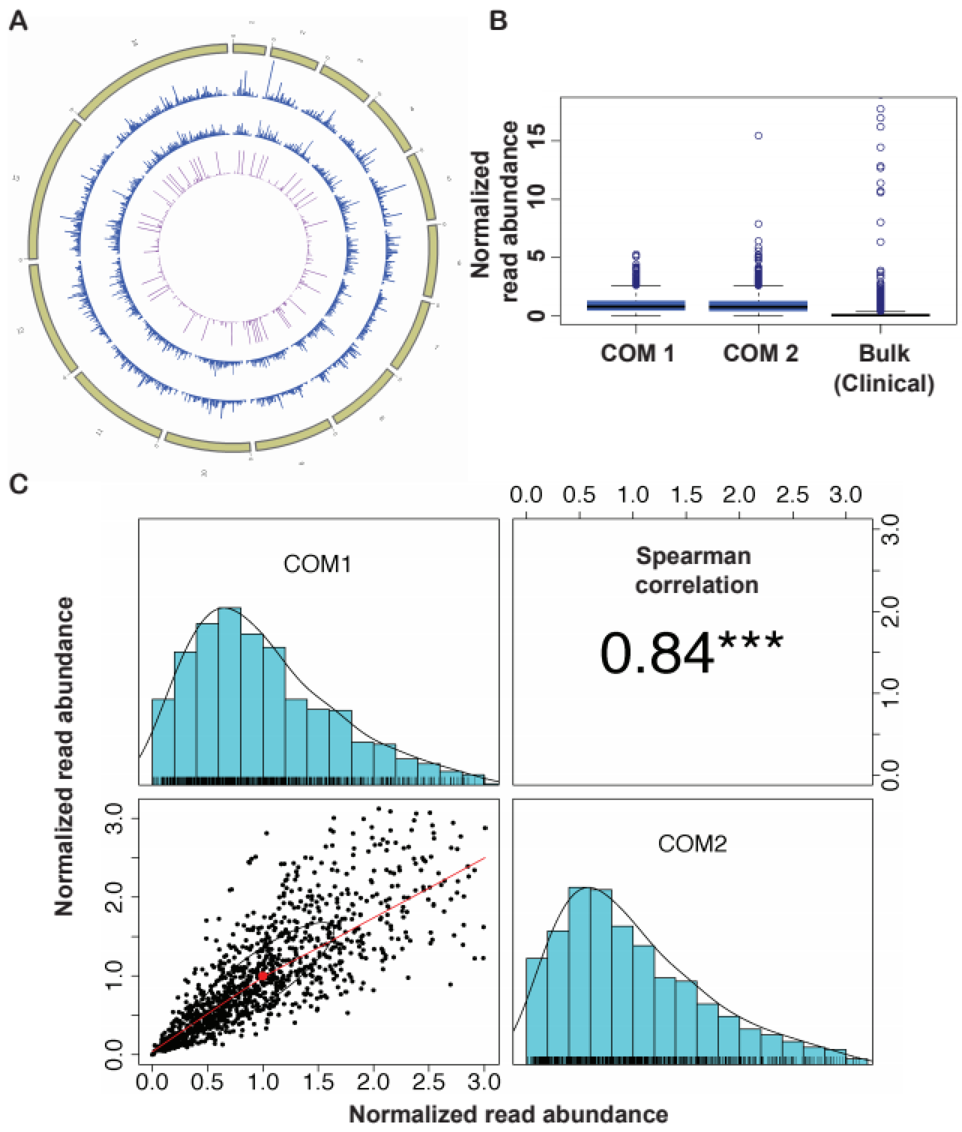


**Figure S4. Uniformity of read abundance across the whole genome and correlation analysis for clinical samples. A. Normalized read abundance across the genome.** Distribution of read abundance in 20kb bins of clinical samples is shown. Read counts in each bin were normalized by the mean read count over the whole genome for each sample. The circles from outside to the inside represent: chromosome 1:14 (green); 2 COM sample (blue); 1 Clinical bulk genomic DNA (pink). **B**. **Distribution of normalized read abundance values for all bins.** Box plot representing the normalized read abundance per 20kb bins, outliers are illustrated by blue dots. **C**. **Paired panels for 1X1 matrices represent Spearman correlation, histogram and pairwise scatterplot among the normalized read abundance (20kb bins) of the COM1 and COM2**

**sample.** Outlier bins were removed, see *Methods* for outlier identification. The Spearman correlation coefficient is listed above the diagonal, and stars indicate the p-value at the levels of 0.1 (no star), 0.05 (*), 0.01 (**), and 0.001 (***). The histograms on the diagonal show the distribution of normalized read abundance in each sample. The scatter plots include a fitted line through the locally smoothed regression and correlation ellipses (an ellipse around the mean with the axis length reflecting one standard deviation of the x and y variables).

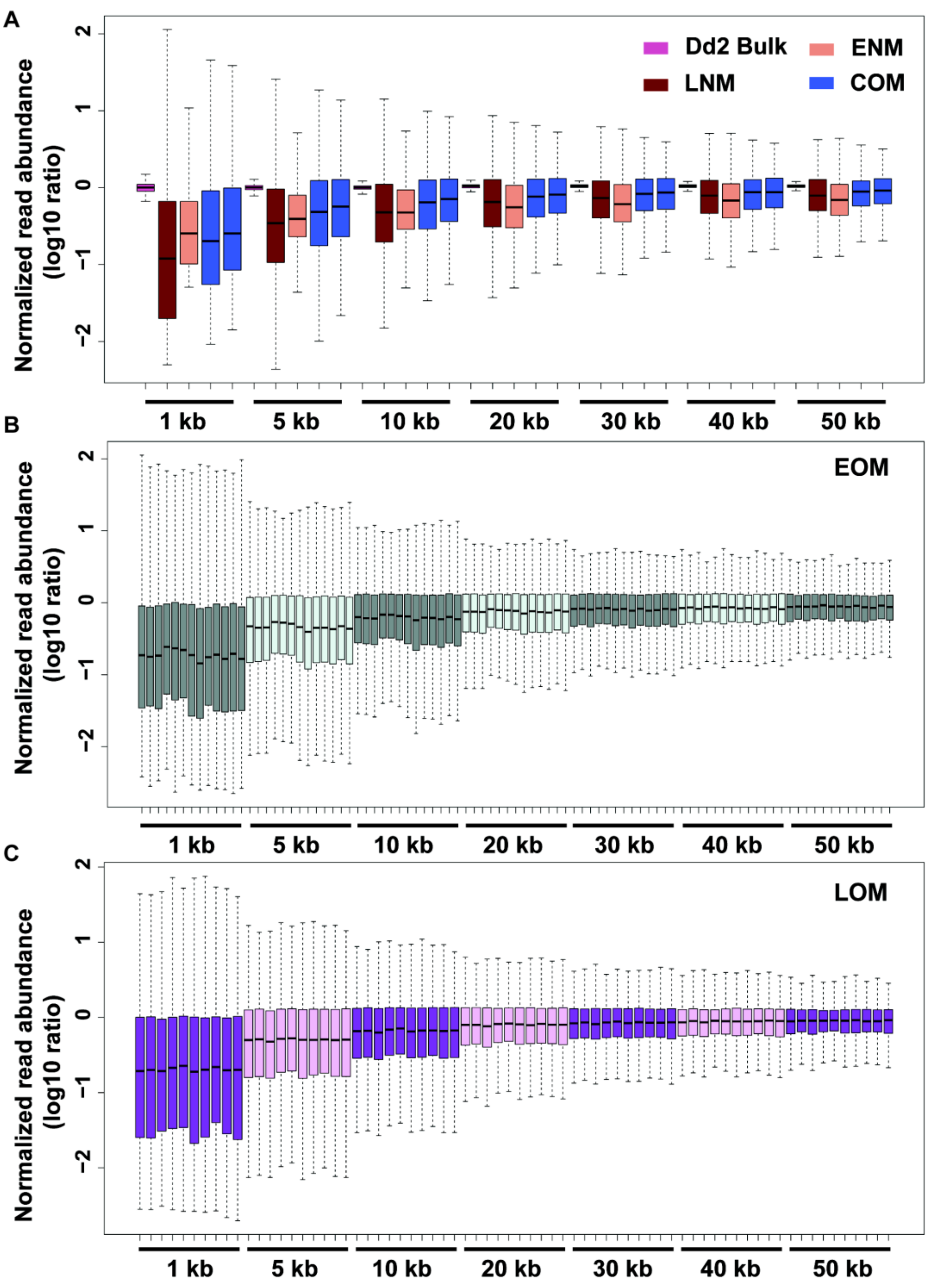


**Figure S5. Distribution of normalized read counts in various bins sizes.** The Log10 ratios of normalized read abundance in 1- 50kb (at intervals of 5 and 10kb) are showed for sequenced samples. The boxes indicate Q1 (25th percentiles) to Q3 (75th percentiles) with a horizontal line drawn in the middle to denote the median. Outliers, above the highest point of the upper whisker (Q3 + 1.5×IQR) or below the lowest point of the lower whisker (Q1-1.5×IQR), are not displayed.

**A. Distribution of normalized read counts in various bins sizes for select sample types.** Dd2 Bulk (purple), ENM (pink, 1 samples), LNM (maroon, 1 samples), COM (blue, 2 samples) samples. **B. Distribution of normalized read counts in various bin sizes for all EOM samples.** EOM (green, n=13). **C. Distribution of normalized read counts in various bin sizes for LOM samples.** LOM (purple, n=10).

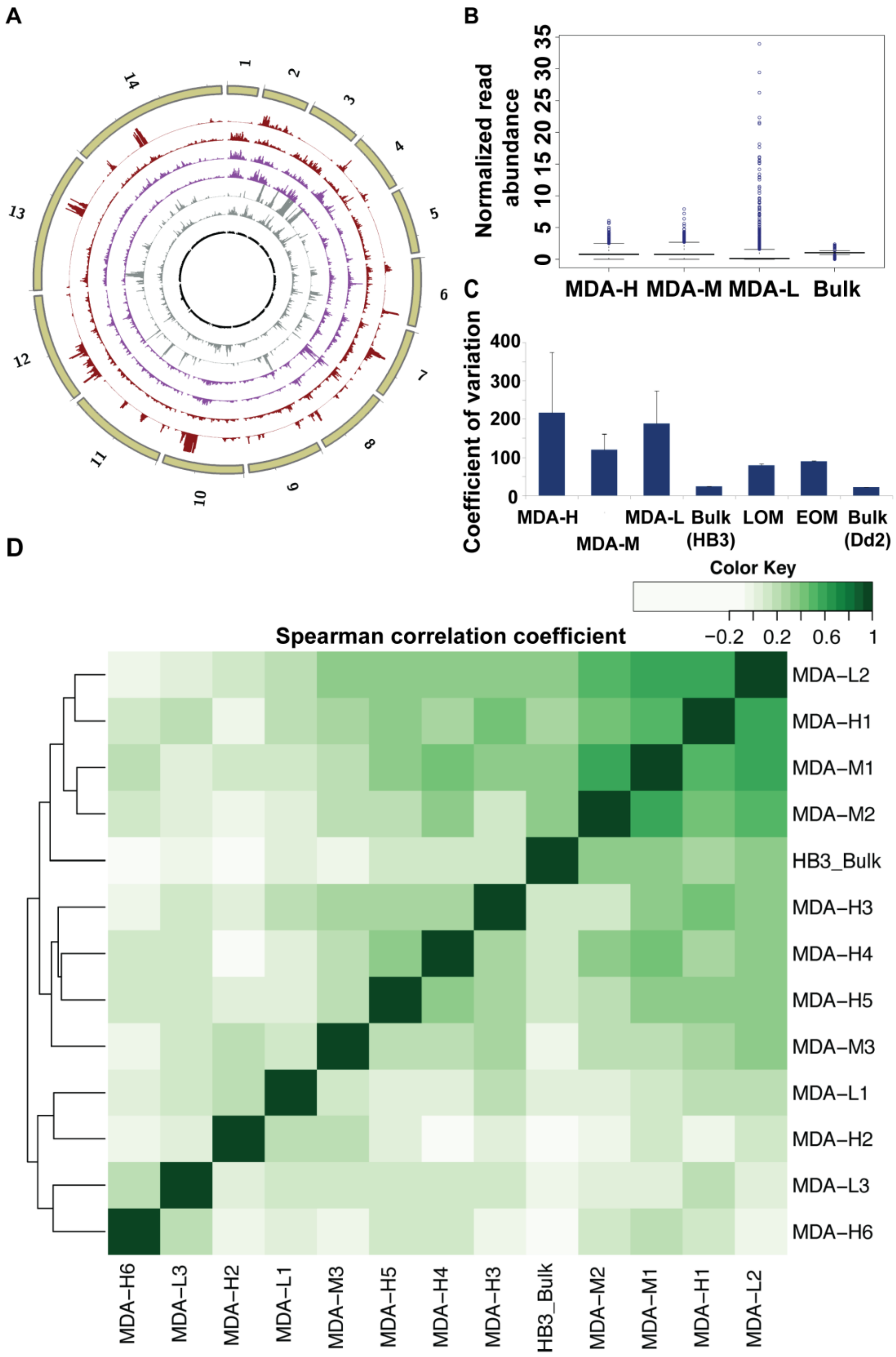


**Figure S6. Uniformity of coverage and correlation analysis in MDA-amplified single cell samples.** Samples for this analysis were amplified and sequence by Trevino et al., 2017 and analyzed using our pipeline (**Figure S2**). **A. Normalized read abundance across the genome.**

Distribution of read abundance in 20kb bins of MDA-amplified samples is shown. Read counts in each bin were normalized by the mean read count over the whole genome for each sample. The circles from outside to the inside represent: chromosome 1:14 (tan); two MDA-H samples (dark red, HB3 parasite with high DNA content amplified by MDA); two MDA-M sample (purple, HB3 parasite with medium DNA content amplified by MDA); two MDA-L samples (grey, parasite with low DNA content amplified by MDA); one HB3 Bulk sample. **B. Distribution of normalized read abundance values for all bins in MDA amplified samples.** MDA-H: 1 representative sample; MDA-M: 1 representative sample; MDA-L: 1 representative sample from (Trevino et al., 2017). **C. Comparison of coefficient of variation of normalized read abundance between MDA and MALBAC amplified single parasites.** The average and SD (error bars) of coefficient of variation of all samples from each type are represented (MDA-H: 6 samples; MDA-M: 3 samples; MDA-L: 3 samples; HB3 Bulk: 1 sample; LOM: 10 samples; EOM: 13 samples; Dd2 Bulk: 1 sample). **D. Spearman correlation coefficient between MDA amplified samples.** The hierarchical clustering heatmap was generated using Spearman correlation coefficients of normalized read abundance. The color scale indicates the degree of correlation (white, low correlation; green, high correlation).
