## Additional file 2 for "Single cell sequencing of the small and AT-skewed genome of malaria parasites"

### Table S1. GC content of 5 base windows in the *P. falciparum* genome

| GC content of 5-base window* | Regions | | | | | |
|---|---|---|---|---|---|---|
| | Whole genome | | Genic regions | | Intergenic regions | |
| | Count of windows | Percentage of windows | Count of windows | Percentage of windows | Count of windows | Percentage of windows |
| 0.00% | 8965130 | 38.49% | 4155040 | 29.87% | 4794912 | 51.34% |
| 20.00% | 8046176 | 34.54% | 5179126 | 37.24% | 2848042 | 30.49% |
| 40.00% | 4613827 | 19.81% | 3311057 | 23.81% | 1294355 | 13.86% |
| 60.00% | 1430717 | 6.14% | 1079231 | 7.76% | 349881 | 3.75% |
| 80.00% | 220354 | 0.95% | 171989 | 1.24% | 48285 | 0.52% |
| 100.00% | 16362 | 0.07% | 11956 | 0.09% | 4399 | 0.05% |

*The analysis of GC content of 5-base window is conducted on the P. falciparum 3D7 genome.

### Table S2. Primer and probe design for droplet digital PCR

| Genes | Gene ID | Locus Name | Chr. | Copy number in Dd2 | Reference | Target | | | |
|---|---|---|---|---|---|---|---|---|---|
| | | | | | | Category | Final concentration | Sequence | Product size |
| Gene-1 | PF3D7_0717700 | *Pfseryl tRNA syn* | 7 | 1 | [7] | F Primer | 900nM | 5'-GGAACAATTCTGTATTGCTTTACC-3' | 142bp |
| | | | | | | R Primer | 900nM | 5'-AAGCTGCGTTGTTTAAAGCTC-3' | |
| | | | | | | Probe | 250nM | 5'-(VIC)-ACATGAAGAAATGATACAAACA-(IOWA BLACK FQ)-3'* | |
| Gene-2 | PF3D7_0818900 | *Pfhsp70* | 8 | 1 | None | F Primer | 600nM | 5'-TGCTGTCATTACCGTTCCAG-3' | 118bp |
| | | | | | | R Primer | 600nM | 5'-AGCAGCTGCAGTAGGTTCATT-3' | |
| | | | | | | Probe | 250nM | 5'-(HEX)-AGATGCTGGTACAATTGCAGGA-(IOWA BLACK FQ)-3'* | |
| Gene-3 | PF3D7_0417200 | *Pfdhfr* | 4 | 1 | None | F Primer | 600nM | 5'-CTCATATGATTGCACAAGTCTG-3' | 133bp |
| | | | | | | R Primer | 600nM | 5'-GTTGGGAATGGATAGGGTATT-3' | |
| | | | | | | Probe | 250nM | 5'-(FAM)-TTGCAACCTGCGCAGTTCATACAC-(IOWA BLACK FQ)-3'* | |
| Gene-4 | PF3D7_1324900 | *Pfldh* | 13 | 1 | [67] | F Primer | 600nM | 5'-ACGATTTGGCTGGAGCAGAT-3' | 85bp |
| | | | | | | R Primer | 600nM | 5'-TCTCTATTCCATTCTTTGTCACTCTTTC-3' | |
| | | | | | | Probe | 250nM | 5'-(HEX)-AGTAATAGTAACAGCTGGATTTACCAAGGCCCCA-(IOWA BLACK FQ)-3'* | |
| Gene-5 | PF3D7_0112300, PF3D7_1148600, PF3D7_1371000 | *Pf18S rDNA* | 1,11, 13 | 3 | [68] | F Primer | 600nM | 5'-CTTTTGAGAGGTTTTGTTACTTTGAGTAA-3' | 99bp |
| | | | | | | R Primer | 600nM | 5'-TATTCCATGCTGTAGTATTCAAACACA-3' | |
| | | | | | | Probe | 250nM | 5'-(FAM)-TGTTCATAACAGACGGGTAGTCATGATTGAGTTCA-(IOWA BLACK FQ)-3'* | |
| Gene-6 | PF3D7_0523000 | *Pfmdr1* | 5 | 3 | None | F Primer | 600nM | 5'-TGCCCACAGAATTGCATCTA-3' | 99bp |
| | | | | | | R Primer | 600nM | 5'-TCGTGTGTTCCATGTGACTG-3' | |
| | | | | | | Probe | 250nM | 5'-(FAM)-ACCCTGATCGAAATGGAACCT-(IOWA BLACK FQ)-3'* | |

*VIC, HEX and FAM, fluorescent reporter dyes; IOWA BLACK FQ: quencher (Integrated DNA Technologies).

**Table S3. Correlation between ddPCR gene copy concentration and sequencing depth of ddPCR target**

| Sample | Gene* | Chr. | ddPCR target | | | | Sequencing read depth of the ddPCR target | Gene copy concentration by ddPCR/20ul reaction | Kendal rank correlation coefficient | Kendal rank correlation P value (two tailed significance test) |
|---|---|---|---|---|---|---|---|---|---|---|
| | | | Start | End | Size of interval | Mappability | | | | |
| EOM 1 | Gene-6 | 5 | 961993 | 962091 | 98 | 1 | 46 | 603 | | |
| EOM 1 | Gene-2 | 8 | 860842 | 860959 | 117 | 1 | 13 | 177 | | |
| EOM 1 | Gene-1 | 7 | 765165 | 765307 | 142 | 1 | 36 | 472 | 1 | 0.014309 |
| EOM 1 | Gene-3 | 4 | 749673 | 749785 | 112 | 1 | 207 | 1131 | | |
| EOM 1 | Gene-4 | 13 | 1041359 | 1041443 | 84 | 1 | 0 | 7.2 | | |
| EOM 1 | Gene-5 | 1 | 474531 | 474629 | 98 | 0.33 | | 22.6 | | |
| EOM 19 | Gene-6 | 5 | 961993 | 962091 | 98 | 1 | 64 | 471 | | |
| EOM 19 | Gene-2 | 8 | 860842 | 860959 | 117 | 1 | 2 | 47.2 | | |
| EOM 19 | Gene-1 | 7 | 765165 | 765307 | 142 | 1 | 38 | 182 | 1 | 0.014309 |
| EOM 19 | Gene-3 | 4 | 749673 | 749785 | 112 | 1 | 369 | 1488 | | |
| EOM 19 | Gene-4 | 13 | 1041359 | 1041443 | 84 | 1 | 0 | 4.1 | | |
| EOM 19 | Gene-5 | 1 | 474531 | 474629 | 98 | 0.33 | | 263 | | |
| EOM 20 | Gene-6 | 5 | 961993 | 962091 | 98 | 1 | 34 | 524 | | |
| EOM 20 | Gene-2 | 8 | 860842 | 860959 | 117 | 1 | 6 | 40.7 | | |
| EOM 20 | Gene-1 | 7 | 765165 | 765307 | 142 | 1 | 36 | 578 | 1 | 0.014309 |
| EOM 20 | Gene-3 | 4 | 749673 | 749785 | 112 | 1 | 126 | 1719 | | |
| EOM 20 | Gene-4 | 13 | 1041359 | 1041443 | 84 | 1 | 0 | 6.5 | | |
| EOM 20 | Gene-5 | 1 | 474531 | 474629 | 98 | 0.33 | | 248 | | |
| EOM 21 | Gene-6 | 5 | 961993 | 962091 | 98 | 1 | 11 | 403 | | |
| EOM 21 | Gene-2 | 8 | 860842 | 860959 | 117 | 1 | 2 | 94 | | |
| EOM 21 | Gene-1 | 7 | 765165 | 765307 | 142 | 1 | 11 | 460 | 0.6 | 0.14167 |
| EOM 21 | Gene-3 | 4 | 749673 | 749785 | 112 | 1 | 67 | 306 | | |
| EOM 21 | Gene-4 | 13 | 1041359 | 1041443 | 84 | 1 | 1 | 2.8 | | |
| EOM 21 | Gene-5 | 1 | 474531 | 474629 | 98 | 0.33 | | 158 | | |
| EOM 22 | Gene-6 | 5 | 961993 | 962091 | 98 | 1 | 41 | 652 | | |
| EOM 22 | Gene-2 | 8 | 860842 | 860959 | 117 | 1 | 9 | 118 | | |
| EOM 22 | Gene-1 | 7 | 765165 | 765307 | 142 | 1 | 17 | 182 | 1 | 0.014309 |
| EOM 22 | Gene-3 | 4 | 749673 | 749785 | 112 | 1 | 311 | 1979 | | |
| EOM 22 | Gene-4 | 13 | 1041359 | 1041443 | 84 | 1 | 0 | 9.9 | | |
| EOM 22 | Gene-5 | 1 | 474531 | 474629 | 98 | 0.33 | | 272 | | |
| EOM 23 | Gene-6 | 5 | 961993 | 962091 | 98 | 1 | 25 | 372 | | |
| EOM 23 | Gene-2 | 8 | 860842 | 860959 | 117 | 1 | 11 | 60.8 | | |
| EOM 23 | Gene-1 | 7 | 765165 | 765307 | 142 | 1 | 8 | 165 | 0.8 | 0.050054 |
| EOM 23 | Gene-3 | 4 | 749673 | 749785 | 112 | 1 | 69 | 427 | | |
| EOM 23 | Gene-4 | 13 | 1041359 | 1041443 | 84 | 1 | 0 | 5.7 | | |
| EOM 23 | Gene-5 | 1 | 474531 | 474629 | 98 | 0.33 | | 74 | | |
| EOM 24 | Gene-6 | 5 | 961993 | 962091 | 98 | 1 | 50 | 817 | | |
| EOM 24 | Gene-2 | 8 | 860842 | 860959 | 117 | 1 | 37 | 977 | | |
| EOM 24 | Gene-1 | 7 | 765165 | 765307 | 142 | 1 | 29 | 989 | 0.4 | 0.327234 |
| EOM 24 | Gene-3 | 4 | 749673 | 749785 | 112 | 1 | 349 | 1324 | | |
| EOM 24 | Gene-4 | 13 | 1041359 | 1041443 | 84 | 1 | 7 | 59.7 | | |
| EOM 24 | Gene-5 | 1 | 474531 | 474629 | 98 | 0.33 | | 513 | | |
| EOM 25 | Gene-6 | 5 | 961993 | 962091 | 98 | 1 | 56 | 452 | | |
| EOM 25 | Gene-2 | 8 | 860842 | 860959 | 117 | 1 | 29 | 375 | | |
| EOM 25 | Gene-1 | 7 | 765165 | 765307 | 142 | 1 | 8 | 103 | 0.4 | 0.327234 |
| EOM 25 | Gene-3 | 4 | 749673 | 749785 | 112 | 1 | 149 | 51.7 | | |
| EOM 25 | Gene-4 | 13 | 1041359 | 1041443 | 84 | 1 | 0 | 13 | | |
| EOM 25 | Gene-5 | 1 | 474531 | 474629 | 98 | 0.33 | | 471 | | |
| EOM 26 | Gene-6 | 5 | 961993 | 962091 | 98 | 1 | 53 | 565 | | |
| EOM 26 | Gene-2 | 8 | 860842 | 860959 | 117 | 1 | 35 | 722 | | |
| EOM 26 | Gene-1 | 7 | 765165 | 765307 | 142 | 1 | 13 | 85 | 0.4 | 0.327234 |
| EOM 26 | Gene-3 | 4 | 749673 | 749785 | 112 | 1 | 15 | 21.4 | | |
| EOM 26 | Gene-4 | 13 | 1041359 | 1041443 | 84 | 1 | 2 | 65 | | |
| EOM 26 | Gene-5 | 1 | 474531 | 474629 | 98 | 0.33 | | 169 | | |
| EOM 27 | Gene-6 | 5 | 961993 | 962091 | 98 | 1 | 50 | 1018 | | |
| EOM 27 | Gene-2 | 8 | 860842 | 860959 | 117 | 1 | 3 | 110 | | |
| EOM 27 | Gene-1 | 7 | 765165 | 765307 | 142 | 1 | 87 | 899 | 0.4 | 0.327234 |
| EOM 27 | Gene-3 | 4 | 749673 | 749785 | 112 | 1 | 141 | 145 | | |
| EOM 27 | Gene-4 | 13 | 1041359 | 1041443 | 84 | 1 | 1 | 40.3 | | |
| EOM 27 | Gene-5 | 1 | 474531 | 474629 | 98 | 0.33 | | 1279 | | |

*Gene 5 (*Pf18S rRNA*) was excluded from correlation analysis due to non-unique reads mapping

**Table S3. Correlation between ddPCR gene copy concentration and sequencing depth of ddPCR target**

| Sample | Gene* | Chr. | ddPCR target | | | | Sequencing read depth of the ddPCR target | Gene copy concentration by ddPCR/20ul reaction | Kendal rank correlation coefficient | Kendal rank correlation P value (two tailed significance test) |
|---|---|---|---|---|---|---|---|---|---|---|
| | | | Start | End | Size of interval | Mappability | | | | |
| EOM 28 | Gene-6 | 5 | 961993 | 962091 | 98 | 1 | 57 | 1008 | | |
| EOM 28 | Gene-2 | 8 | 860842 | 860959 | 117 | 1 | 18 | 480 | | |
| EOM 28 | Gene-1 | 7 | 765165 | 765307 | 142 | 1 | 18 | 56.2 | 0.8 | 0.050054 |
| EOM 28 | Gene-3 | 4 | 749673 | 749785 | 112 | 1 | 206 | 510 | | |
| EOM 28 | Gene-4 | 13 | 1041359 | 1041443 | 84 | 1 | 1 | 46.2 | | |
| EOM 28 | Gene-5 | 1 | 474531 | 474629 | 98 | 0.33 | | 307 | | |
| EOM 29 | Gene-6 | 5 | 961993 | 962091 | 98 | 1 | 36 | 626 | | |
| EOM 29 | Gene-2 | 8 | 860842 | 860959 | 117 | 1 | 24 | 454 | | |
| EOM 29 | Gene-1 | 7 | 765165 | 765307 | 142 | 1 | 25 | 433 | 0.8 | 0.050054 |
| EOM 29 | Gene-3 | 4 | 749673 | 749785 | 112 | 1 | 369 | 1238 | | |
| EOM 29 | Gene-4 | 13 | 1041359 | 1041443 | 84 | 1 | 4 | 62.5 | | |
| EOM 29 | Gene-5 | 1 | 474531 | 474629 | 98 | 0.33 | | 467 | | |
| EOM 30 | Gene-6 | 5 | 961993 | 962091 | 98 | 1 | 77 | 383 | | |
| EOM 30 | Gene-2 | 8 | 860842 | 860959 | 117 | 1 | 37 | 618 | | |
| EOM 30 | Gene-1 | 7 | 765165 | 765307 | 142 | 1 | 5 | 134 | 0.4 | 0.327234 |
| EOM 30 | Gene-3 | 4 | 749673 | 749785 | 112 | 1 | 181 | 136 | | |
| EOM 30 | Gene-4 | 13 | 1041359 | 1041443 | 84 | 1 | 2 | 35.8 | | |
| EOM 30 | Gene-5 | 1 | 474531 | 474629 | 98 | 0.33 | | 222 | | |
| LOM 4 | Gene-6 | 5 | 961993 | 962091 | 98 | 1 | 17 | 582 | | |
| LOM 4 | Gene-2 | 8 | 860842 | 860959 | 117 | 1 | 9 | 363 | | |
| LOM 4 | Gene-1 | 7 | 765165 | 765307 | 142 | 1 | 26 | 813 | 1 | 0.014309 |
| LOM 4 | Gene-3 | 4 | 749673 | 749785 | 112 | 1 | 172 | 2680 | | |
| LOM 4 | Gene-4 | 13 | 1041359 | 1041443 | 84 | 1 | 1 | 26.3 | | |
| LOM 4 | Gene-5 | 1 | 474531 | 474629 | 98 | 0.33 | | 1336 | | |
| LOM 5 | Gene-6 | 5 | 961993 | 962091 | 98 | 1 | 68 | 1359 | | |
| LOM 5 | Gene-2 | 8 | 860842 | 860959 | 117 | 1 | 30 | 543 | | |
| LOM 5 | Gene-1 | 7 | 765165 | 765307 | 142 | 1 | 27 | 1765 | 0.6 | 0.14167 |
| LOM 5 | Gene-3 | 4 | 749673 | 749785 | 112 | 1 | 355 | 3370 | | |
| LOM 5 | Gene-4 | 13 | 1041359 | 1041443 | 84 | 1 | 0 | 44.9 | | |
| LOM 5 | Gene-5 | 1 | 474531 | 474629 | 98 | 0.33 | | 506 | | |
| LOM 11 | Gene-6 | 5 | 961993 | 962091 | 98 | 1 | 39 | 998 | | |
| LOM 11 | Gene-2 | 8 | 860842 | 860959 | 117 | 1 | 60 | 1213 | | |
| LOM 11 | Gene-1 | 7 | 765165 | 765307 | 142 | 1 | 22 | 522 | 1 | 0.014309 |
| LOM 11 | Gene-3 | 4 | 749673 | 749785 | 112 | 1 | 260 | 3510 | | |
| LOM 11 | Gene-4 | 13 | 1041359 | 1041443 | 84 | 1 | 3 | 38.8 | | |
| LOM 11 | Gene-5 | 1 | 474531 | 474629 | 98 | 0.33 | | 1160 | | |
| LOM 12 | Gene-6 | 5 | 961993 | 962091 | 98 | 1 | 55 | 1813 | | |
| LOM 12 | Gene-2 | 8 | 860842 | 860959 | 117 | 1 | 25 | 698 | | |
| LOM 12 | Gene-1 | 7 | 765165 | 765307 | 142 | 1 | 29 | 975 | 1 | 0.014309 |
| LOM 12 | Gene-3 | 4 | 749673 | 749785 | 112 | 1 | 393 | 2900 | | |
| LOM 12 | Gene-4 | 13 | 1041359 | 1041443 | 84 | 1 | 2 | 41.9 | | |
| LOM 12 | Gene-5 | 1 | 474531 | 474629 | 98 | 0.33 | | 1328 | | |
| LOM 13 | Gene-6 | 5 | 961993 | 962091 | 98 | 1 | 99 | 2960 | | |
| LOM 13 | Gene-2 | 8 | 860842 | 860959 | 117 | 1 | 17 | 548 | | |
| LOM 13 | Gene-1 | 7 | 765165 | 765307 | 142 | 1 | 9 | 82 | 0.6 | 0.14167 |
| LOM 13 | Gene-3 | 4 | 749673 | 749785 | 112 | 1 | 237 | 1213 | | |
| LOM 13 | Gene-4 | 13 | 1041359 | 1041443 | 84 | 1 | 2 | 104 | | |
| LOM 13 | Gene-5 | 1 | 474531 | 474629 | 98 | 0.33 | | 642 | | |
| LOM 14 | Gene-6 | 5 | 961993 | 962091 | 98 | 1 | 48 | 642 | | |
| LOM 14 | Gene-2 | 8 | 860842 | 860959 | 117 | 1 | 10 | 251 | | |
| LOM 14 | Gene-1 | 7 | 765165 | 765307 | 142 | 1 | 27 | 201 | 0.8 | 0.050054 |
| LOM 14 | Gene-3 | 4 | 749673 | 749785 | 112 | 1 | 247 | 2126 | | |
| LOM 14 | Gene-4 | 13 | 1041359 | 1041443 | 84 | 1 | 2 | 108 | | |
| LOM 14 | Gene-5 | 1 | 474531 | 474629 | 98 | 0.33 | | 1314 | | |

*Gene 5 (*Pf18S rRNA*) was excluded from correlation analysis due to non-unique reads mapping

**Table S3. Correlation between ddPCR gene copy concentration and sequencing depth of ddPCR target**

| Sample | Gene* | Chr. | ddPCR target | | | | Sequencing read depth of the ddPCR target | Gene copy concentration by ddPCR/20ul reaction | Kendal rank correlation coefficient | Kendal rank correlation P value (two tailed significance test) |
|---|---|---|---|---|---|---|---|---|---|---|
| | | | Start | End | Size of interval | Mappability | | | | |
| LOM 15 | Gene-6 | 5 | 961993 | 962091 | 98 | 1 | 16 | 740 | | |
| LOM 15 | Gene-2 | 8 | 860842 | 860959 | 117 | 1 | 39 | 1103 | | |
| LOM 15 | Gene-1 | 7 | 765165 | 765307 | 142 | 1 | 74 | 922 | 0.8 | 0.050054 |
| LOM 15 | Gene-3 | 4 | 749673 | 749785 | 112 | 1 | 238 | 1172 | | |
| LOM 15 | Gene-4 | 13 | 1041359 | 1041443 | 84 | 1 | 3 | 94 | | |
| LOM 15 | Gene-5 | 1 | 474531 | 474629 | 98 | 0.33 | | 1183 | | |
| LOM 16 | Gene-6 | 5 | 961993 | 962091 | 98 | 1 | 49 | 865 | | |
| LOM 16 | Gene-2 | 8 | 860842 | 860959 | 117 | 1 | 24 | 881 | | |
| LOM 16 | Gene-1 | 7 | 765165 | 765307 | 142 | 1 | 57 | 1189 | 0 | 0 |
| LOM 16 | Gene-3 | 4 | 749673 | 749785 | 112 | 1 | 148 | 686 | | |
| LOM 16 | Gene-4 | 13 | 1041359 | 1041443 | 84 | 1 | 1 | 83 | | |
| LOM 16 | Gene-5 | 1 | 474531 | 474629 | 98 | 0.33 | | 873 | | |
| LOM 17 | Gene-6 | 5 | 961993 | 962091 | 98 | 1 | 86 | 2360 | | |
| LOM 17 | Gene-2 | 8 | 860842 | 860959 | 117 | 1 | 49 | 1291 | | |
| LOM 17 | Gene-1 | 7 | 765165 | 765307 | 142 | 1 | 58 | 1663 | 0.2 | 0.624275 |
| LOM 17 | Gene-3 | 4 | 749673 | 749785 | 112 | 1 | 154 | 349 | | |
| LOM 17 | Gene-4 | 13 | 1041359 | 1041443 | 84 | 1 | 2 | 374 | | |
| LOM 17 | Gene-5 | 1 | 474531 | 474629 | 98 | 0.33 | | 1648 | | |
| LOM 18 | Gene-6 | 5 | 961993 | 962091 | 98 | 1 | 93 | 897 | | |
| LOM 18 | Gene-2 | 8 | 860842 | 860959 | 117 | 1 | 23 | 707 | | |
| LOM 18 | Gene-1 | 7 | 765165 | 765307 | 142 | 1 | 57 | 463 | 0.2 | 0.624275 |
| LOM 18 | Gene-3 | 4 | 749673 | 749785 | 112 | 1 | 124 | 212 | | |
| LOM 18 | Gene-4 | 13 | 1041359 | 1041443 | 84 | 1 | 12 | 182 | | |
| LOM 18 | Gene-5 | 1 | 474531 | 474629 | 98 | 0.33 | | 1532 | | |
| LNM 2 | Gene-6 | 5 | 961993 | 962091 | 98 | 1 | 101 | 770 | | |
| LNM 2 | Gene-2 | 8 | 860842 | 860959 | 117 | 1 | 5 | 325 | | |
| LNM 2 | Gene-1 | 7 | 765165 | 765307 | 142 | 1 | 65 | 10000 | 0.6 | 0.14167 |
| LNM 2 | Gene-3 | 4 | 749673 | 749785 | 112 | 1 | 94 | 759 | | |
| LNM 2 | Gene-4 | 13 | 1041359 | 1041443 | 84 | 1 | 2 | 8.8 | | |
| LNM 2 | Gene-5 | 1 | 474531 | 474629 | 98 | 0.33 | | 5170 | | |
| ENM 3 | Gene-6 | 5 | 961993 | 962091 | 98 | 1 | 9 | 143 | | |
| ENM 3 | Gene-2 | 8 | 860842 | 860959 | 117 | 1 | 0 | 21.6 | | |
| ENM 3 | Gene-1 | 7 | 765165 | 765307 | 142 | 1 | 6 | 6140 | 0.4 | 0.327234 |
| ENM 3 | Gene-3 | 4 | 749673 | 749785 | 112 | 1 | 0 | 6.7 | | |
| ENM 3 | Gene-4 | 13 | 1041359 | 1041443 | 84 | 1 | 0 | 0.4 | | |
| ENM 3 | Gene-5 | 1 | 474531 | 474629 | 98 | 0.33 | | 991 | | |
| Bulk | Gene-6 | 5 | 961993 | 962091 | 98 | 1 | 200 | 61 | | |
| Bulk | Gene-2 | 8 | 860842 | 860959 | 117 | 1 | 130 | 23 | | |
| Bulk | Gene-1 | 7 | 765165 | 765307 | 142 | 1 | 163 | 22 | 0.2 | 0.624275 |
| Bulk | Gene-3 | 4 | 749673 | 749785 | 112 | 1 | 144 | 24 | | |
| Bulk | Gene-4 | 13 | 1041359 | 1041443 | 84 | 1 | 130 | 23 | | |
| Bulk | Gene-5 | 1 | 474531 | 474629 | 98 | 0.33 | | 76 | | |
| COM 1 | Gene-6 | 5 | 961993 | 962091 | 98 | 1 | 4 | 43.7 | | |
| COM 1 | Gene-2 | 8 | 860842 | 860959 | 117 | 1 | 2 | 56.7 | | |
| COM 1 | Gene-1 | 7 | 765165 | 765307 | 142 | 1 | 1 | 7.9 | 0.4 | 0.327234 |
| COM 1 | Gene-3 | 4 | 749673 | 749785 | 112 | 1 | 18 | 37 | | |
| COM 1 | Gene-4 | 13 | 1041359 | 1041443 | 84 | 1 | 0 | 0 | | |
| COM 1 | Gene-5 | 1 | 474531 | 474629 | 98 | 0.33 | | 185 | | |
| COM 2 | Gene-6 | 5 | 961993 | 962091 | 98 | 1 | 8 | 188 | | |
| COM 2 | Gene-2 | 8 | 860842 | 860959 | 117 | 1 | 9 | 145 | | |
| COM 2 | Gene-1 | 7 | 765165 | 765307 | 142 | 1 | 4 | 59 | 0.8 | 0.050054 |
| COM 2 | Gene-3 | 4 | 749673 | 749785 | 112 | 1 | 68 | 314 | | |
| COM 2 | Gene-4 | 13 | 1041359 | 1041443 | 84 | 1 | 0 | 1.9 | | |
| COM 2 | Gene-5 | 1 | 474531 | 474629 | 98 | 0.33 | | 55.6 | | |

*Gene 5 (*Pf18S rRNA*) was excluded from correlation analysis due to non-unique reads mapping

**Table S4. Overall sequencing read output and percentage of aligned reads to *P. falciparum* genome**

| Sample name | Total reads | Reads aligned (Q>=10, non-duplicated) | % Reads aligned (Q>=10, non-duplicated) | Mean coverage (X) | SD coverage (X) | Coverage breadth % |
|---|---|---|---|---|---|---|
| Dd2 Bulk | 14088518 | 12451854 | 88.38% | 75.8308 | 39.7742 | 96.12% |
| Clinical Bulk | 1798478 | 31179 | 1.73% | 0.0345 | 4.137 | 0.3% |
| COM 1 | 3703368 | 1348000 | 36.40% | 7.4389 | 23.5745 | 47.03% |
| COM 2 | 5199958 | 2079068 | 39.98% | 11.6436 | 40.9011 | 48.87% |
| ENM | 3605934 | 330429 | 9.16% | 1.4713 | 10.6785 | 23.04% |
| LNM | 5441240 | 3843328 | 70.63% | 20.4289 | 113.9654 | 47.43% |
| LOM 4 | 7716822 | 6981687 | 90.47% | 39.5272 | 112.0864 | 57.42% |
| LOM 5 | 7960574 | 7162154 | 89.97% | 40.3766 | 130.3619 | 57.64% |
| LOM 11 | 8435214 | 6572607 | 77.92% | 36.7722 | 185.5418 | 53.56% |
| LOM 12 | 8247324 | 7125641 | 86.40% | 40.1058 | 121.7324 | 58.25% |
| LOM 13 | 8440624 | 7491252 | 88.75% | 41.9726 | 128.0252 | 59.33% |
| LOM 14 | 8355392 | 7499920 | 89.76% | 42.309 | 138.4362 | 56.23% |
| LOM 15 | 8755272 | 7713286 | 88.10% | 42.8325 | 135.5189 | 55.89% |
| LOM 16 | 9144356 | 7401206 | 80.94% | 41.2309 | 137.0799 | 57.71% |
| LOM 17 | 10547490 | 9107689 | 86.35% | 50.2924 | 180.1848 | 58.09% |
| LOM 18 | 10855318 | 9879263 | 91.01% | 55.5314 | 153.4196 | 59.07% |
| EOM 1 | 5879596 | 5134985 | 87.34% | 29.045 | 94.6642 | 57.12% |
| EOM 19 | 8522966 | 6990566 | 82.02% | 37.4556 | 121.5732 | 56.71% |
| EOM 20 | 6916038 | 5821742 | 84.18% | 32.6576 | 102.4475 | 56.62% |
| EOM 21 | 4381554 | 3989410 | 91.05% | 21.9652 | 74.3972 | 58.59% |
| EOM 22 | 9211442 | 8441554 | 91.64% | 45.0949 | 133.0433 | 63.56% |
| EOM 23 | 5557586 | 5004172 | 90.04% | 26.8804 | 87.347 | 58.99% |
| EOM 24 | 8002608 | 6636807 | 82.93% | 37.5702 | 109.667 | 56.46% |
| EOM 25 | 9315768 | 7830782 | 84.06% | 43.9508 | 145.9258 | 55.65% |
| EOM 26 | 8492108 | 6808137 | 80.17% | 37.754 | 132.5401 | 56.76% |
| EOM 27 | 9241720 | 7656595 | 82.85% | 42.7219 | 135.2933 | 57.20% |
| EOM 28 | 9166292 | 7767520 | 84.74% | 42.704 | 136.5738 | 56.78% |
| EOM 29 | 10415486 | 8719933 | 83.72% | 49.1051 | 138.0999 | 59.68% |
| EOM 30 | 8883286 | 7431044 | 83.65% | 41.1568 | 131.6227 | 58.06% |
| HB3 Bulk | 17,719,090 | 16,189,815 | 91.37% | 67.2147 | 48.8778 | 95.51% |
| HB3_L_single_cell | 12,245,390 | 10,488,358 | 85.65% | 41.7289 | 183.9036 | 66.40% |
| HB3_L_single_cell | 12,278,504 | 11,389,251 | 92.76% | 45.9091 | 69.8633 | 94.14% |
| HB3_L_single_cell | 18,664,566 | 17,344,347 | 92.93% | 70.812 | 139.5796 | 82.10% |
| HB3_M_single_cell | 11,392,932 | 10,499,255 | 92.16% | 42.6567 | 50.188 | 95.19% |
| HB3_M_single_cell | 6,880,990 | 6,253,026 | 90.87% | 25.2398 | 52.9048 | 86.70% |
| HB3_M_single_cell | 6084642 | 5510823 | 90.57% | 22.3326 | 30.0916 | 93.64% |
| HB3_H_single_cell | 3,895,054 | 3,397,493 | 87.23% | 13.8257 | 78.6302 | 47.35% |
| HB3_H_single_cell | 11,111,292 | 9,920,995 | 89.29% | 40.2235 | 73.7899 | 91.95% |
| HB3_H_single_cell | 3,658,764 | 3,198,863 | 87.43% | 13.0053 | 20.4314 | 84.00% |
| HB3_H_single_cell | 8,534,604 | 7,957,119 | 93.23% | 32.1219 | 125.2137 | 55.52% |
| HB3_H_single_cell | 5,473,840 | 4,919,047 | 89.86% | 20.0882 | 22.2623 | 94.20% |
| HB3_H_single_cell | 9,999,412 | 8,744,829 | 87.45% | 35.4627 | 58.1737 | 92.44% |

**Table S5. Proportion of reads that contain the MALBAC primer common sequence and their alignment to specific genomic regions**

| Sample name | % aligned reads containing MALBAC primer along the genome | % aligned reads with MALBAC primer overlapped with genomic regions | |
|---|---|---|---|
| | | Genic regions | Intergenic regions |
| Dd2 Bulk | 0.04% | 92.99% | 7.01% |
| COM 1 | 37.72% | 90.05% | 9.95% |
| COM 2 | 35.39% | 92.69% | 7.31% |
| ENM | 23.43% | 84.91% | 15.09% |
| LNM | 25.87% | 94.24% | 5.76% |
| EOM 21 | 39.23% | 93.79% | 6.21% |
| EOM 23 | 38.48% | 95.19% | 4.81% |
| LOM 16 | 27.85% | 95.41% | 4.59% |
| LOM 18 | 27.60% | 95.79% | 4.21% |

**Table S6. CNVs detected by LUMPY in all samples (single cell and DD2 bulk)**
Due to the large size of the table, please access via the link: https://github.com/jessicaliu80/Single-cell-sequencing-data.git

**Table S7. CNVs detected by Gingko in all samples (single cell and Dd2 bulk)**
Due to the large size of the table, please access via the link: https://github.com/jessicaliu80/Single-cell-sequencing-data.git

**Table S8. The number of single cell samples processed at each analysis step**

| Steps | Number of samples | | | | |
|---|---|---|---|---|---|
| | ENM | EOM | LNM | LOM | COM |
| Cells amplified by MALBAC | 3 | 42 | 4 | 20 | 4 |
| Cells yielded product | 3 | 18 | 4 | 18 | 4 |
| Cells evaluated by ddPCR | 3 | 17 | 4 | 14 | 4 |
| Cells passed ddPCR quality control (all 6 genes detected) | 1 | 16 | 2 | 12 | 1 |
| Cells sequenced | 1 | 13 | 1 | 10 | 2 |

### Table S9. DNA yield after MALBAC amplification

| Sample type | DNA yield after MALBAC and purification (ng) | | | | | | | | | | |
|---|---|---|---|---|---|---|---|---|---|---|---|
| | Sequencing sample name | EOM | Sequencing sample name | ENM | Sequencing sample name | LOM | Sequencing sample name | LNM | Sequencing sample name | COM | NTC (opt) | NTC (Non-opt) |
| Successfully amplification | | **1408*** | ENM 3 | **1420** | LOM 14 | **278** | LNM 2 | **1480** | | **1368** | Undetectable | 1250 |
| | EOM 25 | **1296** | | **1740** | LOM 18 | **268** | | **1220** | | **1624** | Undetectable | - |
| | EOM 28 | **1264** | | **2120** | | **241** | | **2425** | COM 2 | **347.2** | Undetectable | - |
| | EOM 30 | **992** | | - | LOM 12 | **239** | | **2265** | COM 1 | **84.8** | Undetectable | - |
| | EOM 26 | **776** | | - | | **232** | | - | | - | - | - |
| | EOM 19 | **568** | | - | LOM 16 | **204** | | - | | - | - | - |
| | EOM 29 | **494** | | - | LOM 5 | **199** | | - | | - | - | - |
| | EOM 1 | **456** | | - | LOM 17 | **178** | | - | | - | - | - |
| | EOM 20 | **456** | | - | | **172** | | - | | - | - | - |
| | EOM 27 | **436** | | - | LOM 4 | **165** | | - | | - | - | - |
| | EOM 24 | **418** | | - | LOM 13 | **144** | | - | | - | - | - |
| | | **417** | | - | LOM 11 | **143** | | - | | - | - | - |
| | EOM 22 | **308** | | - | | **142** | | - | | - | - | - |
| | | **152** | | - | LOM 15 | **107** | | - | | - | - | - |
| | EOM 23 | **117.6** | | - | | 125 | | - | | - | - | - |
| | EOM 21 | **79.6** | | - | | 119 | | - | | - | - | - |
| | | **52.4** | | - | | 118 | | - | | - | - | - |
| | | 164 | | - | | 105 | | - | | - | - | - |
| Unsuccessful amplification | | Undetectable (24 samples) | | Undetectable (0 samples) | | Undetectable (2 samples) | | Undetectable (0 samples) | | - | - | - |
| Success rate | | 43% | | 100.00% | | 90% | | 100.00% | | - | - | - |

*Bolded samples were processed for ddPCR evaluation

**Table S10. Primary data from ddPCR detection: calculation of uniformity score and *Pfmdr1* copy number**

| Sample type | Selected for sequencing | ddPCR input / ng | Gene copy concentration by ddPCR / 20ul reaction | | | | | | MDR1 CNV | Locus representation | | | | | | Uniformity score (30 is perfect) |
|---|---|---|---|---|---|---|---|---|---|---|---|---|---|---|---|---|
| | | | MDR1 | HSP70 | SERYL | 18S | DHFR | LDH | | MDR1 | HSP70 | SERYL | 18S | DHFR | LDH | |
| 3D7 Bulk | Y | 0.025 | 61 | 22.8 | 22 | 76 | 24 | 23 | 2.67 | 100.00% | 100.00% | 100.00% | 100.00% | 100.00% | 100.00% | 30 |
| ENM 3 | Y | 1.5 | 143 | 21.6 | 6140 | 991 | 6.7 | 0 | 0.20 | 3.91% | 1.58% | 465.15% | 21.73% | 0.47% | 0.00% | - |
| ENM | | 1.5 | 213 | 234 | 2068 | 1E+06 | 983 | 37.7 | 0.00 | 5.82% | 17.11% | 156.67% | ### | 68.26% | 2.73% | 13,689 |
| ENM | | 1.5 | 0 | 0 | 0 | 0 | 0 | 0 | - | 0.00% | 0.00% | 0.00% | 0.00% | 0.00% | 0.00% | - |
| LNM 2 | Y | 1.5 | 770 | 325 | 100000 | 5170 | 759 | 8.8 | 0.07 | 21.04% | 23.76% | 7575.76% | 113.38% | 52.71% | 0.64% | 13,121 |
| LNM | | 1.5 | 43.3 | 49.7 | 1847 | 4390 | 141 | 0 | 0.07 | 1.18% | 3.63% | 139.92% | 96.27% | 9.79% | 0.00% | - |
| LNM | | 1.5 | 16 | 292 | 1847 | 11000 | 545 | 0 | 0.01 | 0.44% | 21.35% | 139.92% | 241.23% | 37.85% | 0.00% | - |
| LNM | | 1.5 | 89 | 2.9 | 2660 | 285 | 90 | 7.5 | 0.28 | 2.43% | 0.21% | 201.52% | 6.25% | 6.25% | 0.54% | 1,578 |
| COM 1 | Y | 1.5 | 43.7 | 56.7 | 7.9 | 185 | 37 | 0 | 1.32 | 1.19% | 4.14% | 0.60% | 4.06% | 2.57% | 0.00% | - |
| COM 2 | Y | 1.5 | 188 | 145 | 59 | 55.6 | 314 | 1.9 | 2.46 | 5.14% | 10.60% | 4.47% | 1.22% | 21.81% | 0.14% | 369 |
| COM | | 1.5 | 1.9 | 67 | 9.7 | 57.7 | 349 | 0 | 0.04 | 0.05% | 4.90% | 0.73% | 1.27% | 24.24% | 0.00% | - |
| COM | | 1.5 | 124 | 6 | 3.3 | 0 | 0 | 0 | 9.30 | 3.39% | 0.44% | 0.25% | 0.00% | 0.00% | 0.00% | - |
| EOM 19 | Y | 1.5 | 471 | 47.2 | 182 | 263 | 1488 | 4.1 | 1.92 | 12.87% | 3.45% | 13.79% | 5.77% | 103.33% | 0.30% | 550 |
| EOM 25 | Y | 1.5 | 452 | 375 | 103 | 471 | 51.7 | 13 | 3.08 | 12.35% | 27.41% | 7.80% | 10.33% | 3.59% | 0.94% | 99 |
| EOM 26 | Y | 1.5 | 565 | 722 | 85 | 169 | 21.4 | 65 | 3.47 | 15.44% | 52.78% | 6.44% | 3.71% | 1.49% | 4.71% | 112 |
| EOM 30 | Y | 1.5 | 383 | 618 | 134 | 222 | 136 | 35.8 | 2.51 | 10.46% | 45.18% | 10.15% | 4.87% | 9.44% | 2.59% | 70 |
| EOM 1 | Y | 1.5 | 603 | 177 | 472 | 22.6 | 1131 | 7.2 | 2.50 | 16.48% | 12.94% | 35.76% | 0.50% | 78.54% | 0.52% | 589 |
| EOM 20 | Y | 1.5 | 524 | 40.7 | 578 | 248 | 1719 | 6.5 | 1.68 | 14.32% | 2.98% | 43.79% | 5.44% | 119.38% | 0.47% | 505 |
| EOM 28 | Y | 1.5 | 1008 | 480 | 56.2 | 307 | 510 | 46.2 | 4.19 | 27.54% | 35.09% | 4.26% | 6.73% | 35.42% | 3.35% | 81 |
| EOM 23 | Y | 1.5 | 372 | 60.8 | 165 | 74 | 427 | 5.7 | 3.37 | 10.16% | 4.44% | 12.50% | 1.62% | 29.65% | 0.41% | 198 |
| EOM 21 | Y | 1.5 | 403 | 94 | 460 | 158 | 306 | 2.8 | 2.83 | 11.01% | 6.87% | 34.85% | 3.46% | 21.25% | 0.20% | 423 |
| EOM 22 | Y | 1.5 | 652 | 118 | 182 | 272 | 1979 | 9.9 | 2.03 | 17.81% | 8.63% | 13.79% | 5.96% | 137.43% | 0.72% | 328 |
| EOM 27 | Y | 1.5 | 1018 | 110 | 899 | 1279 | 145 | 40.3 | 2.92 | 27.81% | 8.04% | 68.11% | 28.05% | 10.07% | 2.92% | 89 |
| EOM 29 | Y | 1.5 | 626 | 454 | 433 | 467 | 1238 | 62.5 | 1.91 | 17.10% | 33.19% | 32.80% | 10.24% | 85.97% | 4.53% | 77 |
| EOM 24 | Y | 1.5 | 817 | 977 | 989 | 513 | 1324 | 59.7 | 1.75 | 22.32% | 71.42% | 74.92% | 11.25% | 91.94% | 4.33% | 105 |
| EOM | | 1.5 | 718 | 424 | 632 | 75 | 785 | 2.6 | 2.72 | 19.62% | 30.99% | 47.88% | 1.64% | 54.51% | 0.19% | 929 |
| EOM | | 1.5 | 2880 | 47 | 0 | 32.9 | 82 | 0 | 9.47 | 78.69% | 3.44% | 0.00% | 0.72% | 5.69% | 0.00% | - |
| EOM | | 1.5 | 266 | 92 | 341 | 66 | 1525 | 2.8 | 1.16 | 7.27% | 6.73% | 25.83% | 1.45% | 105.90% | 0.20% | 871 |
| EOM | | 1.5 | 230 | 1.3 | 74.1 | 67.4 | 1249 | 77 | 1.35 | 6.28% | 0.10% | 5.61% | 1.48% | 86.74% | 5.58% | 1,235 |
| LOM 16 | Y | 1.5 | 865 | 881 | 1189 | 873 | 686 | 83 | 1.89 | 23.63% | 64.40% | 90.08% | 19.14% | 47.64% | 6.01% | 71 |
| LOM 18 | Y | 1.5 | 897 | 707 | 463 | 1532 | 212 | 182 | 2.25 | 24.51% | 51.68% | 35.08% | 33.60% | 14.72% | 13.19% | 39 |
| LOM 17 | Y | 1.5 | 2360 | 1291 | 1663 | 1648 | 349 | 374 | 3.07 | 64.48% | 94.37% | 125.98% | 36.14% | 24.24% | 27.10% | 46 |
| LOM 4 | Y | 1.5 | 582 | 363 | 813 | 1336 | 2680 | 26.3 | 1.00 | 15.90% | 26.54% | 61.59% | 29.30% | 186.11% | 1.91% | 213 |
| LOM 5 | Y | 1.5 | 1359 | 543 | 1765 | 506 | 3370 | 44.9 | 1.79 | 37.13% | 39.69% | 133.71% | 11.10% | 234.03% | 3.25% | 206 |
| LOM 13 | Y | 1.5 | 2960 | 548 | 82 | 642 | 1213 | 104 | 5.33 | 80.87% | 40.06% | 6.21% | 14.08% | 84.24% | 7.54% | 90 |
| LOM 12 | Y | 1.5 | 1813 | 698 | 975 | 1328 | 2900 | 41.9 | 2.34 | 49.54% | 51.02% | 73.86% | 29.12% | 201.39% | 3.04% | 166 |
| LOM 14 | Y | 1.5 | 642 | 251 | 201 | 1314 | 2126 | 108 | 1.38 | 17.54% | 18.35% | 15.23% | 28.82% | 147.64% | 7.83% | 76 |
| LOM 15 | Y | 1.5 | 740 | 1103 | 922 | 1183 | 1172 | 94 | 1.42 | 20.22% | 80.63% | 69.85% | 25.94% | 81.39% | 6.81% | 72 |
| LOM 11 | Y | 1.5 | 998 | 1213 | 522 | 1160 | 3510 | 38.8 | 1.34 | 27.27% | 88.67% | 39.55% | 25.44% | 243.75% | 2.81% | 196 |
| LOM | | 1.5 | 2346 | 858 | 1333 | 1167 | 3360 | 5.5 | 2.59 | 64.10% | 62.72% | 100.98% | 25.59% | 233.33% | 0.40% | 1,258 |
| LOM | | 1.5 | 254 | 988 | 366 | 391 | 253 | 7.4 | 1.12 | 6.94% | 72.22% | 27.73% | 8.57% | 17.57% | 0.54% | 292 |
| LOM | | 1.5 | 971 | 2090 | 1312 | 165 | 828 | 0 | 1.81 | 26.53% | 152.78% | 99.39% | 3.62% | 57.50% | 0.00% | - |
| LOM | | 1.5 | 4040 | 256 | 846 | 273 | 1564 | 0 | 5.79 | 110.38% | 18.71% | 64.09% | 5.99% | 108.61% | 0.00% | - |

**Table S11. Coefficient of variation of normalized read abundance in each sequenced sample**

| Batch | Sample name | Coefficient of Variation (CV) of normalized read abundance (%) | AVG CV in each type of samples (%) | SD of CVs in each type of samples |
|---|---|---|---|---|
| - | Dd2 Bulk | 22 | 22 | - |
| - | ENM | 147 | 147 | - |
| - | EOM 1 | 91 | | |
| | EOM 19 | 86 | | |
| | EOM 20 | 91 | | |
| | EOM 25 | 95 | | |
| Batch-1 | EOM 26 | 91 | | |
| | EOM 28 | 92 | | |
| | EOM 30 | 92 | 89 | 4 |
| | EOM 21 | 82 | | |
| Batch-2 | EOM 22 | 81 | | |
| | EOM 23 | 90 | | |
| | EOM 24 | 87 | | |
| Batch-3 | EOM 27 | 91 | | |
| | EOM 29 | 84 | | |
| - | LNM | 111 | 111 | - |
| | EOM 11 | 84 | | |
| | EOM 12 | 77 | | |
| | EOM 13 | 76 | | |
| | EOM 14 | 80 | | |
| | EOM 15 | 79 | | |
| Same Batch | EOM 16 | 78 | 79 | 2 |
| | EOM 17 | 81 | | |
| | EOM 18 | 79 | | |
| | EOM 4 | 82 | | |
| | EOM 5 | 79 | | |
| Same Batch | COM 1 | 78 | 87 | 12 |
| | COM 2 | 96 | | |
| - | Clinical Bulk | 472 | 472 | - |

**Table S12. The equality of CVs in normalized read abundance for sequenced samples**

| Sample name | Paired test on the equality of CVs between a ENM and a EOM | | Sample name | Paired test on the equality of CVs between a LNM and a LOM | |
|---|---|---|---|---|---|
| | Test statistic | P-value | | Test statistic | P-value |
| EOM 1 | 66.76287 | 3.06E-16 | LOM 11 | 30.05426 | 4.20E-08 |
| EOM 19 | 86.53364 | 1.37E-20 | LOM 12 | 54.05277 | 1.95E-13 |
| EOM 20 | 68.38409 | 1.35E-16 | LOM 13 | 60.58646 | 7.04E-15 |
| EOM 21 | 103.6689 | 2.39E-24 | LOM 14 | 43.05383 | 5.33E-11 |
| EOM 22 | 110.2728 | 8.54E-26 | LOM 15 | 47.97997 | 4.31E-12 |
| EOM 23 | 73.3549 | 1.08E-17 | LOM 16 | 52.48721 | 4.33E-13 |
| EOM 24 | 81.76793 | 1.53E-19 | LOM 17 | 41.60157 | 1.12E-10 |
| EOM 25 | 56.81782 | 4.78E-14 | LOM 18 | 45.69159 | 1.38E-11 |
| EOM 26 | 67.57568 | 2.03E-16 | LOM 4 | 36.88771 | 1.25E-09 |
| EOM 27 | 66.63139 | 3.27E-16 | LOM 5 | 48.01785 | 4.22E-12 |
| EOM 28 | 65.25652 | 6.58E-16 | | | |
| EOM 29 | 97.36433 | 5.77E-23 | | | |
| EOM 30 | 65.9547 | 4.61E-16 | | | |

**Table S13. Spearman correlation coefficient of all sequenced samples**

| Spearman correlation coefficient | LOM 4 | LOM 5 | LOM 11 | LOM 12 | LOM 13 | LOM 14 | LOM 15 | LOM 16 | LOM 17 | LOM 18 | EOM 1 | EOM 19 | EOM 20 | EOM 21 | EOM 22 | EOM 23 | EOM 24 | EOM 25 | EOM 26 | EOM 27 | EOM 28 | EOM 29 | EOM 30 | LNM | ENM | Dd2 Bulk |
|---|---|---|---|---|---|---|---|---|---|---|---|---|---|---|---|---|---|---|---|---|---|---|---|---|---|---|
| **LOM 4** | 1.00 | 0.88 | 0.81 | 0.88 | 0.89 | 0.89 | 0.84 | 0.89 | 0.87 | 0.90 | 0.81 | 0.82 | 0.81 | 0.76 | 0.85 | 0.80 | 0.87 | 0.89 | 0.81 | 0.82 | 0.87 | 0.87 | 0.83 | 0.55 | 0.53 | 0.24 |
| **LOM 5** | 0.88 | 1.00 | 0.83 | 0.90 | 0.91 | 0.91 | 0.87 | 0.91 | 0.90 | 0.91 | 0.82 | 0.85 | 0.83 | 0.78 | 0.88 | 0.83 | 0.89 | 0.90 | 0.86 | 0.85 | 0.89 | 0.89 | 0.85 | 0.53 | 0.51 | 0.26 |
| **LOM 11** | 0.81 | 0.83 | 1.00 | 0.83 | 0.83 | 0.84 | 0.79 | 0.83 | 0.81 | 0.84 | 0.76 | 0.76 | 0.76 | 0.71 | 0.80 | 0.74 | 0.81 | 0.82 | 0.77 | 0.77 | 0.80 | 0.81 | 0.77 | 0.53 | 0.50 | 0.26 |
| **LOM 12** | 0.88 | 0.90 | 0.83 | 1.00 | 0.94 | 0.91 | 0.86 | 0.90 | 0.89 | 0.91 | 0.83 | 0.85 | 0.83 | 0.78 | 0.88 | 0.83 | 0.89 | 0.91 | 0.84 | 0.83 | 0.88 | 0.89 | 0.85 | 0.56 | 0.53 | 0.22 |
| **LOM 13** | 0.89 | 0.91 | 0.83 | 0.94 | 1.00 | 0.92 | 0.87 | 0.91 | 0.90 | 0.92 | 0.84 | 0.85 | 0.84 | 0.80 | 0.89 | 0.85 | 0.90 | 0.90 | 0.85 | 0.85 | 0.89 | 0.90 | 0.86 | 0.54 | 0.52 | 0.26 |
| **LOM 14** | 0.89 | 0.91 | 0.84 | 0.91 | 0.92 | 1.00 | 0.86 | 0.92 | 0.89 | 0.92 | 0.82 | 0.84 | 0.82 | 0.77 | 0.87 | 0.82 | 0.90 | 0.92 | 0.84 | 0.84 | 0.89 | 0.90 | 0.85 | 0.56 | 0.52 | 0.25 |
| **LOM 15** | 0.84 | 0.87 | 0.79 | 0.86 | 0.87 | 0.86 | 1.00 | 0.87 | 0.85 | 0.87 | 0.78 | 0.80 | 0.78 | 0.75 | 0.84 | 0.79 | 0.84 | 0.85 | 0.81 | 0.79 | 0.84 | 0.84 | 0.81 | 0.53 | 0.48 | 0.23 |
| **LOM 16** | 0.89 | 0.91 | 0.83 | 0.90 | 0.91 | 0.92 | 0.87 | 1.00 | 0.90 | 0.91 | 0.82 | 0.84 | 0.84 | 0.78 | 0.87 | 0.82 | 0.89 | 0.90 | 0.85 | 0.85 | 0.89 | 0.89 | 0.86 | 0.55 | 0.52 | 0.25 |
| **LOM 17** | 0.87 | 0.90 | 0.81 | 0.89 | 0.90 | 0.89 | 0.85 | 0.90 | 1.00 | 0.89 | 0.80 | 0.82 | 0.80 | 0.75 | 0.85 | 0.79 | 0.87 | 0.89 | 0.82 | 0.81 | 0.88 | 0.87 | 0.83 | 0.54 | 0.52 | 0.22 |
| **LOM 18** | 0.90 | 0.91 | 0.84 | 0.91 | 0.92 | 0.92 | 0.87 | 0.91 | 0.89 | 1.00 | 0.84 | 0.84 | 0.83 | 0.79 | 0.89 | 0.83 | 0.90 | 0.91 | 0.84 | 0.86 | 0.88 | 0.91 | 0.86 | 0.55 | 0.54 | 0.28 |
| **EOM 1** | 0.81 | 0.82 | 0.76 | 0.83 | 0.84 | 0.82 | 0.78 | 0.82 | 0.80 | 0.84 | 1.00 | 0.80 | 0.80 | 0.84 | 0.89 | 0.84 | 0.82 | 0.82 | 0.79 | 0.79 | 0.83 | 0.82 | 0.81 | 0.48 | 0.43 | 0.29 |
| **EOM 19** | 0.82 | 0.85 | 0.76 | 0.85 | 0.85 | 0.84 | 0.80 | 0.84 | 0.82 | 0.84 | 0.80 | 1.00 | 0.79 | 0.77 | 0.86 | 0.81 | 0.84 | 0.85 | 0.80 | 0.78 | 0.85 | 0.84 | 0.82 | 0.50 | 0.48 | 0.21 |
| **EOM 20** | 0.81 | 0.83 | 0.76 | 0.83 | 0.84 | 0.82 | 0.78 | 0.84 | 0.80 | 0.83 | 0.80 | 0.79 | 1.00 | 0.79 | 0.86 | 0.80 | 0.81 | 0.82 | 0.79 | 0.76 | 0.83 | 0.82 | 0.80 | 0.50 | 0.47 | 0.26 |
| **EOM 21** | 0.76 | 0.78 | 0.71 | 0.78 | 0.80 | 0.77 | 0.75 | 0.78 | 0.75 | 0.79 | 0.84 | 0.77 | 0.79 | 1.00 | 0.93 | 0.89 | 0.78 | 0.76 | 0.75 | 0.74 | 0.79 | 0.76 | 0.78 | 0.40 | 0.39 | 0.33 |
| **EOM 22** | 0.85 | 0.88 | 0.80 | 0.88 | 0.89 | 0.87 | 0.84 | 0.87 | 0.85 | 0.89 | 0.89 | 0.86 | 0.86 | 0.93 | 1.00 | 0.93 | 0.87 | 0.87 | 0.83 | 0.82 | 0.88 | 0.86 | 0.86 | 0.50 | 0.47 | 0.32 |
| **EOM 23** | 0.80 | 0.83 | 0.74 | 0.83 | 0.85 | 0.82 | 0.79 | 0.82 | 0.79 | 0.83 | 0.84 | 0.81 | 0.80 | 0.89 | 0.93 | 1.00 | 0.82 | 0.82 | 0.79 | 0.79 | 0.82 | 0.81 | 0.81 | 0.46 | 0.42 | 0.30 |
| **EOM 24** | 0.87 | 0.89 | 0.81 | 0.89 | 0.90 | 0.90 | 0.84 | 0.89 | 0.87 | 0.90 | 0.82 | 0.84 | 0.81 | 0.78 | 0.87 | 0.82 | 1.00 | 0.88 | 0.84 | 0.84 | 0.88 | 0.90 | 0.85 | 0.53 | 0.51 | 0.28 |
| **EOM 25** | 0.89 | 0.90 | 0.82 | 0.91 | 0.90 | 0.92 | 0.85 | 0.90 | 0.89 | 0.91 | 0.82 | 0.85 | 0.82 | 0.76 | 0.87 | 0.82 | 0.88 | 1.00 | 0.85 | 0.84 | 0.90 | 0.88 | 0.86 | 0.56 | 0.53 | 0.21 |
| **EOM 26** | 0.81 | 0.86 | 0.77 | 0.84 | 0.85 | 0.84 | 0.81 | 0.85 | 0.82 | 0.84 | 0.79 | 0.80 | 0.79 | 0.75 | 0.83 | 0.79 | 0.84 | 0.85 | 1.00 | 0.78 | 0.84 | 0.82 | 0.79 | 0.50 | 0.47 | 0.22 |
| **EOM 27** | 0.82 | 0.85 | 0.77 | 0.83 | 0.85 | 0.84 | 0.79 | 0.85 | 0.81 | 0.86 | 0.79 | 0.78 | 0.76 | 0.74 | 0.82 | 0.79 | 0.84 | 0.84 | 0.78 | 1.00 | 0.83 | 0.84 | 0.80 | 0.51 | 0.49 | 0.29 |
| **EOM 28** | 0.87 | 0.89 | 0.80 | 0.88 | 0.89 | 0.89 | 0.84 | 0.89 | 0.88 | 0.88 | 0.83 | 0.85 | 0.83 | 0.79 | 0.88 | 0.82 | 0.88 | 0.90 | 0.84 | 0.83 | 1.00 | 0.88 | 0.85 | 0.54 | 0.50 | 0.23 |
| **EOM 29** | 0.87 | 0.89 | 0.81 | 0.89 | 0.90 | 0.90 | 0.84 | 0.89 | 0.87 | 0.91 | 0.82 | 0.84 | 0.82 | 0.76 | 0.86 | 0.81 | 0.90 | 0.88 | 0.82 | 0.84 | 0.88 | 1.00 | 0.85 | 0.56 | 0.53 | 0.26 |
| **EOM 30** | 0.83 | 0.85 | 0.77 | 0.85 | 0.86 | 0.85 | 0.81 | 0.86 | 0.83 | 0.86 | 0.81 | 0.82 | 0.80 | 0.78 | 0.86 | 0.81 | 0.85 | 0.86 | 0.79 | 0.80 | 0.85 | 0.85 | 1.00 | 0.51 | 0.48 | 0.26 |
| **LNM** | 0.55 | 0.53 | 0.53 | 0.56 | 0.54 | 0.56 | 0.53 | 0.55 | 0.54 | 0.55 | 0.48 | 0.50 | 0.50 | 0.40 | 0.50 | 0.46 | 0.53 | 0.56 | 0.50 | 0.51 | 0.54 | 0.56 | 0.51 | 1.00 | 0.73 | 0.07 |
| **ENM** | 0.53 | 0.51 | 0.50 | 0.53 | 0.52 | 0.52 | 0.48 | 0.52 | 0.52 | 0.54 | 0.43 | 0.48 | 0.47 | 0.39 | 0.47 | 0.42 | 0.51 | 0.53 | 0.47 | 0.49 | 0.50 | 0.53 | 0.48 | 0.73 | 1.00 | 0.05 |
| **Dd2 Bulk** | 0.24 | 0.26 | 0.26 | 0.22 | 0.26 | 0.25 | 0.23 | 0.25 | 0.22 | 0.28 | 0.29 | 0.21 | 0.26 | 0.33 | 0.32 | 0.30 | 0.28 | 0.21 | 0.22 | 0.29 | 0.23 | 0.26 | 0.26 | 0.07 | 0.05 | 1.00 |

**Table S14. Comparison between experimental conditions of MALBAC and MDA-amplified single cell samples**

| Steps | Single cell samples | |
|---|---|---|
| | **MALBAC** | **MDA*** |
| Parasite line | Dd2 | HB3 |
| Culture | RPMI +Albumax | RPMI +Albumax |
| Cell staining | SYBR green | Vibrant Dye Cycle Green dye |
| Cell sorting | CellRaft AIR System | Fluorescence-Activated Cell Sorting |
| Cell lysis | -80°C storage and MALBAC lysis | -80°C storage and Repli-g MDA lysis |
| Amplification | Optimized MALBAC | Repli-g MDA |
| Sequencing platform | Illumina Nextseq 550 using 150 bp paired end | Illumina HiSeq 2500 using 101 bp paired end |

*MDA sample experimental conditions are from the Trevino SG et al., 2017 study.

**Table S15. The coefficient of variation of normalized read abundance in MDA amplified samples**

| Sample name | Coefficient of Variation (CV) of normalized read abundance (%) | AVG CV in each type of samples(%) | SD of CVs in each type of samples |
|---|---|---|---|
| HB3 Bulk | 24 | 24 | - |
| MDA-H1 | 83 | 217 | 157 |
| MDA-H2 | 324 | | |
| MDA-H3 | 127 | | |
| MDA-H4 | 134 | | |
| MDA-H5 | 148 | | |
| MDA-H6 | 489 | | |
| MDA-M1 | 103 | 120 | 40 |
| MDA-M2 | 91 | | |
| MDA-M3 | 166 | | |
| MDA-L1 | 166 | 188 | 86 |
| MDA-L2 | 115 | | |
| MDA-L3 | 283 | | |

**Table S16. Coverage comparison after downsampling to 300,000 reads**

| Studies | Sample name | Coverage (X) | GC % | Coverage breadth | | |
|---|---|---|---|---|---|---|
| | | | | Whole genome | Genic regions | Intergenic regions |
| MALBAC | ENM | 1.3263 | 25.17% | 21.71% | 32.93% | 4.99% |
| | LNM | 1.6246 | 24.32% | 26.18% | 40.34% | 5.12% |
| | LOM 4 | 1.7025 | 22.23% | 32.56% | 50.62% | 5.73% |
| | LOM 5 | 1.6953 | 22.37% | 32.35% | 50.21% | 5.80% |
| | LOM 11 | 1.6802 | 22.75% | 30.29% | 46.74% | 5.80% |
| | LOM 12 | 1.6956 | 22.17% | 32.50% | 50.33% | 5.97% |
| | LOM 13 | 1.6867 | 22.25% | 32.91% | 50.90% | 6.15% |
| | LOM 14 | 1.6986 | 22.55% | 32.12% | 50.08% | 5.41% |
| | LOM 15 | 1.6697 | 22.41% | 31.55% | 49.03% | 5.55% |
| | LOM 16 | 1.6716 | 22.51% | 32.42% | 50.21% | 5.94% |
| | LOM 17 | 1.6633 | 22.68% | 31.63% | 49.23% | 5.43% |
| | LOM 18 | 1.6924 | 22.20% | 32.59% | 50.52% | 5.91% |
| | EOM 1 | 1.6971 | 20.79% | 31.30% | 47.29% | 7.49% |
| | EOM 19 | 1.6095 | 21.48% | 29.51% | 45.11% | 6.24% |
| | EOM 20 | 1.691 | 21.35% | 31.13% | 47.51% | 6.75% |
| | EOM 21 | 1.6536 | 19.44% | 31.67% | 46.11% | 10.19% |
| | EOM 22 | 1.6033 | 20.63% | 32.07% | 48.42% | 7.73% |
| | EOM 23 | 1.613 | 20.24% | 31.22% | 46.64% | 8.22% |
| | EOM 24 | 1.7013 | 21.95% | 31.90% | 49.15% | 6.24% |
| | EOM 25 | 1.6843 | 22.39% | 29.41% | 46.20% | 4.45% |
| | EOM 26 | 1.664 | 21.82% | 29.76% | 45.60% | 6.17% |
| | EOM 27 | 1.6763 | 22.22% | 31.30% | 48.36% | 5.94% |
| | EOM 28 | 1.6523 | 21.78% | 29.88% | 46.53% | 5.11% |
| | EOM 29 | 1.693 | 22.05% | 32.32% | 49.70% | 6.49% |
| | EOM 30 | 1.6614 | 21.67% | 30.05% | 46.46% | 5.63% |
| | Dd2 Bulk | 1.8516 | 18.84% | 76.83% | 80.63% | 71.17% |
| | COM 1 | 1.6492 | 20.35% | 31.40% | 46.85% | 8.47% |
| | COM 2 | 1.6783 | 21.14% | 30.72% | 47.08% | 6.42% |
| MDA | MDA-H | 1.2414 | 20.84% | 53.90% | 63.56% | 39.76% |
| | | 1.2264 | 21.52% | 22.72% | 25.74% | 18.31% |
| | | 1.2351 | 20.98% | 45.65% | 53.57% | 34.06% |
| | | 1.2335 | 20.98% | 45.53% | 53.43% | 33.92% |
| | | 1.2328 | 21.12% | 44.86% | 53.48% | 32.23% |
| | | 1.2371 | 21.57% | 19.56% | 23.53% | 13.70% |
| | MDA-M | 1.2304 | 21.29% | 50.64% | 60.94% | 35.52% |
| | | 1.2343 | 21.13% | 51.81% | 61.89% | 37.01% |
| | | 1.2262 | 21.39% | 40.54% | 47.85% | 29.83% |
| | MDA-L | 1.2413 | 21.05% | 37.11% | 43.47% | 27.79% |
| | | 1.2243 | 22.08% | 47.41% | 59.01% | 30.36% |
| | | 1.2058 | 22.19% | 24.47% | 28.36% | 18.72% |
| | HB3 bulk | 1.2529 | 20.96% | 63.35% | 73.09% | 48.86% |

**Table S17. Spearman correlation coefficient of MDA amplified samples**

| Spearman correlation coefficient | MDA-L1 | MDA-L2 | MDA-L3 | MDA-M1 | MDA-M2 | MDA-M3 | MDA-H1 | MDA-H2 | MDA-H3 | MDA-H4 | MDA-H5 | MDA-H6 | HB3 Bulk |
|---|---|---|---|---|---|---|---|---|---|---|---|---|---|
| **MDA-L1** | 1.00 | 0.23 | 0.15 | 0.16 | 0.07 | 0.13 | 0.21 | 0.21 | 0.16 | 0.03 | 0.07 | 0.05 | 0.04 |
| **MDA-L2** | 0.23 | 1.00 | 0.04 | 0.55 | 0.52 | 0.33 | 0.56 | 0.13 | 0.35 | 0.33 | 0.32 | 0.00 | 0.32 |
| **MDA-L3** | 0.15 | 0.04 | 1.00 | 0.03 | 0.03 | 0.10 | 0.18 | 0.02 | 0.12 | 0.15 | 0.14 | 0.17 | -0.04 |
| **MDA-M1** | 0.16 | 0.55 | 0.03 | 1.00 | 0.61 | 0.18 | 0.49 | 0.10 | 0.33 | 0.44 | 0.34 | 0.17 | 0.34 |
| **MDA-M2** | 0.07 | 0.52 | 0.03 | 0.61 | 1.00 | 0.20 | 0.41 | -0.03 | 0.14 | 0.34 | 0.17 | 0.10 | 0.36 |
| **MDA-M3** | 0.13 | 0.33 | 0.10 | 0.18 | 0.20 | 1.00 | 0.27 | 0.18 | 0.26 | 0.19 | 0.19 | -0.01 | -0.02 |
| **MDA-H1** | 0.21 | 0.56 | 0.18 | 0.49 | 0.41 | 0.27 | 1.00 | 0.00 | 0.46 | 0.24 | 0.37 | 0.16 | 0.25 |
| **MDA-H2** | 0.21 | 0.13 | 0.02 | 0.10 | -0.03 | 0.18 | 0.00 | 1.00 | 0.06 | -0.11 | 0.04 | -0.06 | -0.15 |
| **MDA-H3** | 0.16 | 0.35 | 0.12 | 0.33 | 0.14 | 0.26 | 0.46 | 0.06 | 1.00 | 0.29 | 0.24 | -0.02 | 0.13 |
| **MDA-H4** | 0.03 | 0.33 | 0.15 | 0.44 | 0.34 | 0.19 | 0.24 | -0.11 | 0.29 | 1.00 | 0.31 | 0.09 | 0.14 |
| **MDA-H5** | 0.07 | 0.32 | 0.14 | 0.34 | 0.17 | 0.19 | 0.37 | 0.04 | 0.24 | 0.31 | 1.00 | 0.10 | 0.14 |
| **MDA-H6** | 0.05 | 0.00 | 0.17 | 0.17 | 0.10 | -0.01 | 0.16 | -0.06 | -0.02 | 0.09 | 0.10 | 1.00 | -0.09 |
| **HB3 Bulk** | 0.04 | 0.32 | -0.04 | 0.34 | 0.36 | -0.02 | 0.25 | -0.15 | 0.13 | 0.14 | 0.14 | -0.09 | 1.00 |

**Table S18. Detection of true CNVs in single cell genomes by discordant/split read or read depth analysis**

| Detection method | CNV name | Bin size | No. of single cells with CNV / No. of single cell samples | | | |
|---|---|---|---|---|---|---|
| | | | ENM | EOM | LNM | LOM |
| Lumpy | *Pfmdr1* | - | 0/1 | 2/13 | 0/1 | 1/10 |
| | *Pf11-1* | - | 1/1 | 11/13 | 0/1 | 9/10 |
| | *Pf332* | - | 0/1 | 0/13 | 0/1 | 0/10 |
| Ginkgo | *Pfmdr1* | 1kb | 1/1 | 13/13 | 1/1 | 10/10 |
| | | 5kb | 0/1 | 6/13 | 0/1 | 2/10 |
| | | 8kb | 0/1 | 1/13 | 0/1 | 1/10 |
| | | 10kb | 0/1 | 1/13 | 0/1 | 0/10 |
| | *Pf11-1* | 1kb | 1/1 | 3/13 | 0/1 | 1/10 |
| | | 5kb | 0/1 | 0/13 | 0/1 | 0/10 |
| | | 8kb | N/A | N/A | N/A | N/A |
| | | 10kb | N/A | N/A | N/A | N/A |
| | *Pf332* | 1kb | 1/1 | 1/13 | 0/1 | 0/10 |
| | | 5kb | N/A | N/A | N/A | N/A |
| | | 8kb | N/A | N/A | N/A | N/A |
| | | 10kb | N/A | N/A | N/A | N/A |

Any overlap with the 3 CNVs is considered as detected.

**Table S19. Single cell CNVs detected by both discordant/split read and read depth analysis (excluding true CNVs presented in Table 5)**

| Sample name | Chr. | Start Position | Size (bp) | Type | Support read | | Start Position | Size (bp) | Copy number detected by Ginkgo in different bin sizes | | | | Overlapped size (bp) | Overlapped proportion in Lumpy detected CNV | Overlapped proportion in Ginkgo detected CNV |
|---|---|---|---|---|---|---|---|---|---|---|---|---|---|---|---|
| | | | | | Discordant read | Split read | | | 1kb | 5kb | 8kb | 10kb | | | |
| LNM 2 | 13 | 1979452 | 66800 | DUP | 0 | 2 | 1986000 | 62000 | 3 | - | - | - | 60252 | 90% | 97% |
| ENM 3 | 5 | 375286 | 9588 | DUP | 1 | 1 | 372000 | 16000 | 6 | N/A | N/A | N/A | 9588 | 100% | 60% |
| LOM 4 | 4 | 759338 | 32187 | DUP | 0 | 2 | 751000 | 25000 | 6 | - | N/A | N/A | 16662 | 52% | 67% |
| LOM 12 | 11 | 200087 | 103546 | DEL | 1 | 1 | 174000 | 94000 | 0 | - | - | - | 67913 | 66% | 72% |
| LOM 18 | 10 | 530064 | 42365 | DUP | 0 | 2 | 510000 | 84000 | 2 | - | - | - | 42365 | 100% | 50% |
| EOM 19 | 14 | 1717516 | 21447 | DUP | 1 | 1 | 1716000 | 33000 | 7 | - | N/A | N/A | 21447 | 100% | 65% |
| EOM 19 | 5 | 250670 | 14616 | DEL | 28 | 13 | 251000 | 14000 | 0 | N/A | N/A | N/A | 14000 | 96% | 100% |
| EOM 20 | 1 | 397645 | 11557 | DUP | 26 | 34 | 396000 | 16000 | 6 | N/A | N/A | N/A | 11557 | 100% | 72% |
| EOM 20 | 3 | 705702 | 3550 | DUP | 5 | 4 | 703000 | 7000 | 12 | N/A | N/A | N/A | 3550 | 100% | 51% |
| EOM 20 | 6 | 790445 | 19729 | DUP | 17 | 8 | 790000 | 38000 | 2 | - | N/A | N/A | 19729 | 100% | 52% |
| EOM 20 | 8 | 274934 | 20790 | DEL | 1 | 1 | 278000 | 14000 | 0 | N/A | N/A | N/A | 14000 | 67% | 100% |
| EOM 23 | 6 | 1266293 | 12486 | DUP | 1 | 2 | 1262000 | 17000 | 5 | N/A | N/A | N/A | 12486 | 100% | 73% |
| EOM 23 | 7 | 544707 | 47283 | DUP | 2 | 0 | 544000 | 30000 | 5 | - | N/A | N/A | 29293 | 62% | 98% |
| EOM 24 | 11 | 562230 | 5730 | DUP | 1 | 1 | 561000 | 9000 | 4 | N/A | N/A | N/A | 5730 | 100% | 64% |
| EOM 26 | 12 | 1418656 | 5918 | DUP | 1 | 2 | 1417000 | 9000 | 16 | N/A | N/A | N/A | 5918 | 100% | 66% |
| EOM 27 | 14 | 654902 | 80858 | DUP | 1 | 1 | 694000 | 48000 | 4 | - | - | N/A | 41760 | 52% | 87% |
| EOM 28 | 10 | 884189 | 22283 | DUP | 1 | 1 | 876000 | 24000 | 10 | N/A | N/A | N/A | 15811 | 71% | 66% |
| EOM 29 | 11 | 786733 | 104177 | DUP | 1 | 1 | 776000 | 63000 | 3 | - | - | - | 52267 | 50% | 83% |
| EOM 29 | 11 | 856172 | 54794 | DUP | 1 | 2 | 873000 | 71000 | 3 | - | - | - | 37966 | 69% | 53% |
| EOM 30 | 13 | 1285335 | 81344 | DEL | 1 | 1 | 1233000 | 114000 | 0 | - | - | - | 61665 | 76% | 54% |
